# Identification of Immunological Features Enables Survival Prediction of Muscle-Invasive Bladder Cancer Patients Using Machine Learning

**DOI:** 10.1101/2020.02.24.963181

**Authors:** Christos G Gavriel, Neofytos Dimitriou, Nicolas Brieu, Ines P Nearchou, Ognjen Arandjelović, Günter Schmidt, David J Harrison, Peter D Caie

## Abstract

Clinical staging and prognosis of muscle-invasive bladder cancer (MIBC) routinely includes assessment of patient tissue samples by a pathologist. Recent studies corroborate the importance of image analysis in identifying and quantifying immunological markers from tissue samples that can provide further insights into patient prognosis. In this paper, we apply multiplex immunofluorescence on MIBC tissue sections to capture whole slide images and quantify potential prognostic markers related to lymphocytes, macrophages, tumour buds, and PD-L1. We propose a machine learning based approach for the prediction of 5 year prognosis with different combinations of image, clinical, and spatial features. An ensemble model comprising several functionally different models successfully stratifies MIBC patients into two risk groups with high statistical significance (*p* value < 1*e* − 05). Critical to improving MIBC survival rates, our method classifies correctly 71.4% of the patients who succumb to MIBC within 5 years, significantly higher than the 28.6% of the current clinical gold standard, the TNM staging system.

## Introduction

Urothelial cancer of the bladder (bladder cancer) is one of the most prevalent cancers worldwide with approximately 430,000 new diagnoses each year [1]. High morbidity and mortality rates as well as high socioeconomic burden make bladder cancer a debilitating and often fatal disease [2, 3]. Even though the majority of bladder cancer patients are diagnosed with non-muscle-invasive bladder cancer (NMIBC), recurrence and progression of the disease may lead to muscle-invasive bladder cancer (MIBC) [4]. Approximately 25% of newly diagnosed patients have MIBC, and unlike NMIBC, these tumours are biologically aggressive with limited therapeutic options. Although radical cystectomy with bilateral pelvic lymph node dissection is the current gold-standard treatment for MIBC, more than 50% of MIBC patients die from metastatic disease within 5 years [5]. To decrease the mortality rates, patients with high risk of disease-specific death need to be identified more precisely, thereby allowing better patient management and new treatments to be tested in the high risk group.

Due to the intra- and inter-tumoural heterogeneity of MIBC, evident from the phenotypic and molecular diversity of tumour cells, choosing the most effective treatment for each patient is very challenging [6, 7]. Currently, clinical assessment of bladder uses the Tumour-Node-Metastasis (TNM) staging system [8, 9], where T describes the depth of invasion into the bladder wall, and N and M the presence or lack of node and distant metastasis, respectively. MIBC ranges from tumours which invade the detrusor muscle (T2), to tumours which spread to nearby organs (T4) [10, 1]. Although TNM has a critical role in guiding treatment planning, it remains an anatomy-based classification tool with patients of the same tumour stage experiencing a high variability in disease outcome [11, 12, 13, 14].

The tumour mass comprises a heterogeneous population of cancer cells, and together with a diverse group of resident and infiltrating host cells, they make up the tumour-immune microenvironment [15, 16, 17]. With the emergence of immuno-oncology, many cancer researchers have investigated the importance of the intra-tumoural host immune response within what is often an immunosuppressive tumour microenvironment [18, 19]. Analysing the location, density, functional state, and organisation of the immune cell populations within the tumour landscape, often termed as the immune contexture, has become a fundamental step in identifying immune system characteristics that may be beneficial to patients [20]. In particular, an increasing number of studies has shown the critical role of the immune contexture on patient survivability, suggesting that it could be a valuable determinant of patient prognosis [21, 19]. Motivated by multiple papers [22, 23, 24, 25], we investigate the prognostic role of tumour infiltrating lymphocytes (TILs), tumour-associated macrophages (TAMs), tumour buds (TBs), and programmed cell death-ligand 1 (PD-L1) in MIBC patients.

Lymphocytes and macrophages are generally found either infiltrating into or surrounding the tumour mass, both the core and the invasive front. TILs can be divided into subpopulations by virtue of their specialised functions, surface cluster of differentiation (CD) molecules and, in certain circumstances, morphological features. Cytotoxic T-cells are the main effector cells in anti-tumour T-cell response, with a large volume of studies showing that their presence in the tumour-immune microenvironment is strongly associated with prolonged survival in various types of cancers [26, 27]. TAMs have also been identified as decisive factors in orchestrating the tumour-immune microenvironment [28, 29]. They can exhibit polarised phenotypes, with classically activated M1 and alternatively activated M2 subpopulations possessing anti-tumoural and pro-tumoural capabilities, respectively [30]. In particular, during metastasis, M2 macrophages are recruited at distinct pre-metastatic niches where they can promote tumour cell dissemination and disrupt the function of TILs [28].

Tumour budding is generally considered to be the first step of cancer metastasis, defined as the dissociation of isolated single cancer cells or discrete clusters of up to four cancer cells predominantly from the invasive front of the tumour [31]. Over the last decade, tumour budding has been widely investigated as a marker of aggressive tumour behaviour due to its association with adverse clinicopathological characteristics and with the epithelial-mesenchymal transition [32, 33, 34]. As a result, tumour budding has been added to TNM as a supplementary prognostic factor for colorectal cancer [35]. However, without reliable quantitative methods, tumour budding quantification becomes challenging due to poor inter-observer consistency. [36, 37].

Immune checkpoints are cell surface receptors expressed by immune cells that modulate immune responses [38]. Complex interactions between the immune system and cancer, including manipulation of immune checkpoints such as programmed cell death 1 (PD1), enable tumour cells to evade immune surveillance. Specifically, PD-L1, which is secreted by tumour cells, binds to PD-1 expressed on the surface of TILs and suppresses their function to ensure the growth and development of tumour cells [39]. Although our understanding of the intricate and dynamic relationship between tumour cells and host cells is increasing, further characterization of the precise impact of tumour cells on their surroundings is needed.

In recent years, machine learning (ML) methodologies have been widely utilized for constructing predictive models based on biological features [40, 41, 42]. Nevertheless, the number of papers which have adopted ML methodologies for survival analysis is limited [40, 42, 43, 44]. In this paper, we employ a ML methodology to investigate the role of TILs, TAMs, TBs, PD-L1, and other clinicopathological factors in MIBC patient prognosis. The contribution of this paper is fourfold. To the best of our knowledge, this is the first study reporting the labelling of entire MIBC tissue sections with multiple fluorescence immune markers as well as the quantification of immunological features across both the tumour core and the invasive front of whole slide immunofluorescence images. In addition, using image, spatial, and clinical features, our ML methodology improves the accuracy of 5 year prognosis for MIBC patients by a large margin when compared to the current gold standard, TNM. And lastly, our findings reinforce the importance of the immune contexture in cancer prognosis, thereby further supporting its adoption into the clinic.

## Results

### Patient characteristics

A total of 78 patients diagnosed with MIBC were included in this study. Median age of the patients was 68 years (range 29–87 years) with 43 males and 35 females. According to the TNM staging system guidelines [9], our cohort consists of 17 stage II, 29 stage IIIA, 5 stage IIIB, and 27 stage IV patients. Twenty seven patients had distant metastasis at time of surgery. No positive lymph nodes were found in 57 patients and 1–2 lymph nodes contained tumour cells in 21 patients. Of the 78 patients, 53 patients died due to bladder cancer. Follow-up information was available for all the patients (range 1–113 months). The clinicopathological characteristics of the cohort are summarised in Table 1.

**Table 1:** Patient cohort characteristics.

| Characteristics | Summary |
| --- | --- |
| **MIBC patients** | $N = 78$ |
| **Median survival** (range) | 19 (1-113) months |
| **Age** | $66 \pm 11$ years |
| **Gender** | 55% Male; 45% Female |
| **TNM stage** |  |
| II | 17 (22%) |
| IIIA | 29 (37%) |
| IIIB | 5 (6%) |
| IV | 27 (35%) |
| **Tumour (T)** |  |
| T2 | 18 (23%) |
| T3 | 39 (50%) |
| T4 | 21 (27%) |
| **Node (N)** |  |
| N0 | 57 (73%) |
| N1 | 13 (17%) |
| N2 | 8 (10%) |
| **Metastasis (M)** |  |
| M0 | 51 (65%) |
| M1 | 27 (35%) |

### Fully automated feature extraction

The entire formalin-fixed paraffin-embedded (FFPE) tissue section of each MIBC patient was digitized into a WSI, encompassing both muscle-invasive urothelial carcinoma as well as adjacent benign tissue. Multiplex immunofluorescence, using tyramide signal amplification (TSA), enabled the detection of TILs (general CD3 and cytotoxic CD8 T-cells), TAMs (total CD68 macrophages and M2 CD163 macrophages), PD-L1^+^ cells, cell nuclei (Hoechst), and epithelial cancer cells (Pancytokeratin) including TBs across the WSI of each patient. Machine learning-based image analysis allowed for the localization of each cell, subsequently classified depending on its immunofluorescence (IF) signal as either a: 1) TB, 2) M1 macrophage, 3) M2 macrophage, 4) total macrophage, 5) general T cell, 6) cytotoxic T cell, or 7) PD-L1^+^ cell. Based on the above seven classes, a total of 186 quantitative features were extracted from the tumour core and invasive front of each WSI including the number and density of different cell types, the total size of tumour areas, as well as the pairwise spatial distributions between immune and cancer cells. The tumour core is defined as the main tumour mass and the invasive front as the border of the tumour core with a width of 1000*µm* (500*µm* inside and 500*µm* outside of the border defining the invasive frontin and frontout, respectively) as shown in Supplementary Figure S2. Feature extraction was performed using Definiens Tissue Phenomics® software (Definiens AG, Munich, Germany) [45, 46, 47]. A list of all extracted features is provided as Supplementary Information.

### Feature space and feature selection

In order to capture multiple aspects of the disease, features from both clinical reports and whole slide immunofluorescence images were quantified. Herein, the number and density of PD-L1 positive and negative immune cell populations, as well as of TBs from the WSIs are labelled as “image features”. The pairwise spatial distributions between immune cells and TBs are termed “spatial features”. And lastly, clinicopathological features such as age, gender, and TNM stage are termed “clinical features”. Altogether 201 features were quantified – 126 image, 60 spatial, and 15 clinical features (the complete feature list is provided as Supplementary Information). To investigate whether smaller feature spaces result in better ML models, we ran the same ML workflow over different feature sets. In particular, our experiments were based on the following 7 feature sets: (i) image, (ii) spatial, (iii) clinical, image and spatial, (v) image and clinical, (vi) spatial and clinical, (vii) image, spatial, and clinical.

### Machine learning models and optimizing metric

Five ML algorithms with different theoretical underpinnings were selected to investigate whether the extracted features could predict 5 year survivability in MIBC patients; decision tree (DT), random forest (RF), support vector machine (SVM), logistic regression (LR), and *k* nearest neighbours (KNN). The optimizing metric throughout experimentation was the area under the receiver operating characteristic (AUROC). At the final evaluation phase, classification accuracy, sensitivity, specificity, F1 score, and hazard ratios were also computed for ease of comparative analysis, as shown in Table 2. In order to compute the aforementioned metrics, optimal threshold values were automatically selected at the final stage based on the training set performance. Hazard ratios and the confidence intervals were calculated using univariate Cox regression.

**Table 2:** Comparison between our ensemble model and TNM staging. * 95% Confidence Interval.

| Evaluation metrics | Training set |  | Testing set |  |
| --- | --- | --- | --- | --- |
|  | Ensemble model | TNM staging | Ensemble model | TNM staging |
| AUROC | **98.3** | 71.6 | **89.3** | 64.3 |
| Accuracy | **94.8** | 65.5 | **80.0** | 50.0 |
| Sensitivity | **89.5** | **89.5** | **100.0** | **100.0** |
| Specificity | **97.4** | 53.8 | **71.4** | 28.6 |
| F1 score | **83.3** | 67.7 | **83.3** | 44.0 |
| Hazard ratio | **45.9**<br>(6.2, 341.1)* | 4.4<br>(2.3, 8.6)* | **32.5**<br>(3.9, 270.3)* | 3.3<br>(1.0, 11.0)* |

### Proposed ensemble model

Nested cross validation was implemented to avoid overfitting while maximizing the predictive performance of ML classifiers. In addition, a separate test set was held aside to estimate the generalization performance of the final classifier.

For each of the tested feature sets, the classifier with the highest average AUROC was selected. In case of similar average AUROC between two ML classifiers using the same feature set, we selected the one exhibiting the least variance. The results are shown in Supplementary Table S3. Since multiple classifiers exhibited similar performance across the different feature sets, instead of employing a single classifier, we combined the best ones into an ensemble model. In particular, our ensemble model consists of a LSVM that uses image features (72.80 ± 0.30 AUROC), a DT that uses image and clinical features (68.80 ± 0.80 AUROC), a LR that uses image and spatial features (70.20 ± 14.70 AUROC), and a RF that uses all features (67.30 ± 5.80 AUROC). Following hyperparameter tuning for each one of the selected classifiers on the whole training set, without cross validation, our ensemble model was evaluated on the independent testing set achieving 89.3% AUROC and a highly significant separation of patients into low and high risk groups (*p* value= 7*e* − 06). Patients were classified as high risk by the ensemble model if two or more of the submodels predicted a bad prognosis.

### Pessimistic bias

The large difference between the generalization estimates of algorithm selection and performance evaluation (see Supplementary Table S3 and Table 2) can be mostly attributed to pessimistic bias [48]. Given an already small data set, withholding half of the training data set for evaluation, due to twofold cross validation, increases the chance that a ML model will underfit, i.e. its maximum representation capacity will not be reached [48]. Therefore, the generalization estimate from performance evaluation (Table 2) is more reliable since the whole training set was used.

### Comparing against TNM staging

In order to compare against the gold standard in clinical practice, TNM, patients had to be stratified into low and high risk groups. Based on a pairwise log-rank test comparison in the training data set (results shown in Supplementary Tables S8, S9, and S10), stage II and III patients were considered as the low risk group whereas stage IV patients were considered as the high risk group. The Kaplan-Meier and ROC curves of TNM staging and our ensemble model on the testing set are shown in Figure 1. To allow further comparative analysis, Kaplan-Meier curves of other clinicopathological features such as age and gender are shown in Supplementary Figure S4. In addition, the Kaplan-Meier and ROC curves of each submodel of the ensemble model are shown in Supplementary Figure S5.

**Figure 1:**
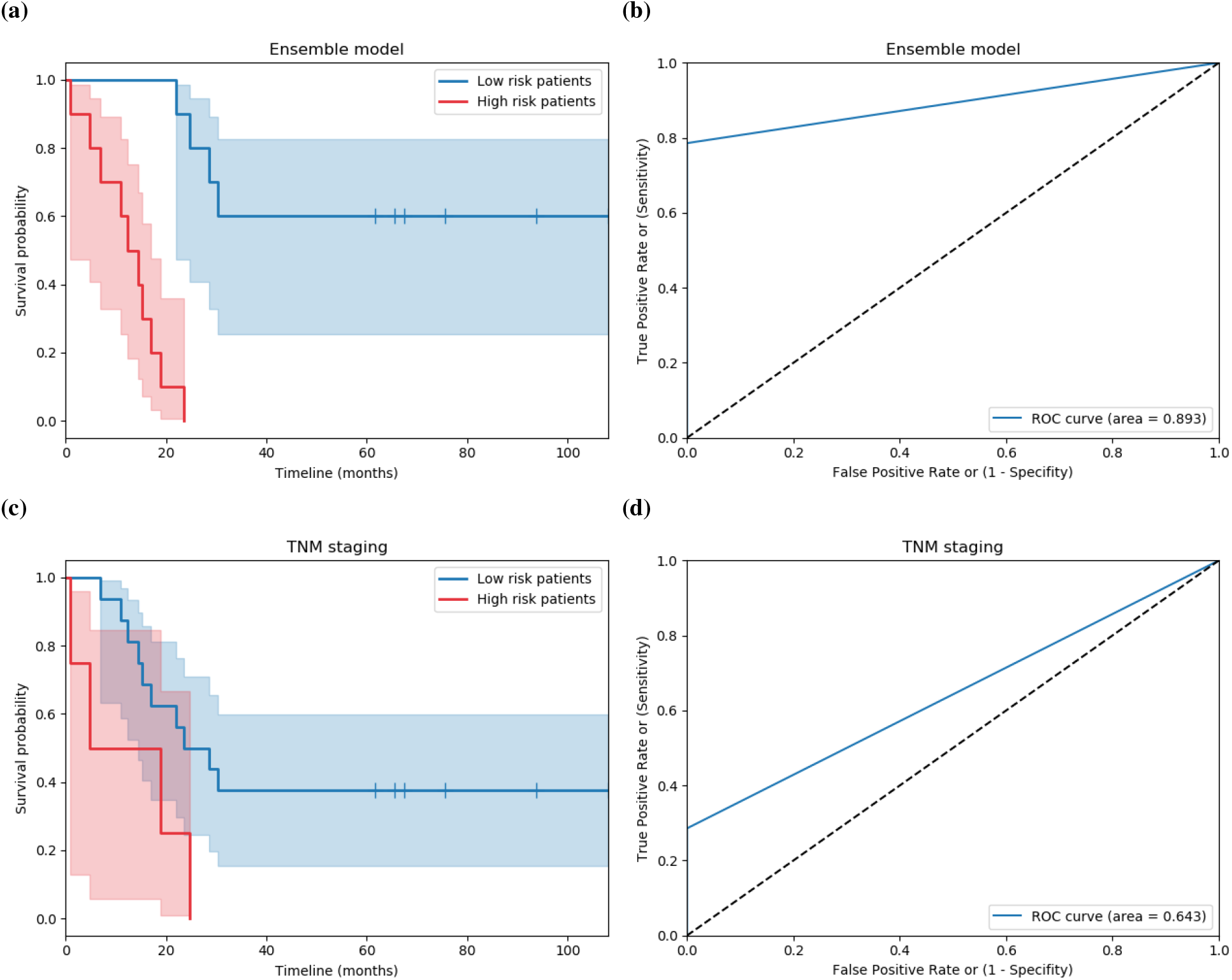
Kaplan-Meier and ROC curves on the testing set for our ensemble model and TNM. Separation was significant based on the (a) ensemble model (*p* value = 7*e* − 06, *N*_*LowRisk*_ = 10 & *N*_*HighRisk*_ = 10) and (c) TNM (*p* value = 0.04, *N*_*LowRisk*_ = 16 & *N*_*HighRisk*_ = 4).

### Post-hoc analysis of features

For each classifier of the ensemble model, post-hoc analysis was conducted to reveal the features guiding survivability prediction. The feature considered at each node of a DT is readily interpretable (see Supplementary Figure S8). For the LR, its coefficients determine the importance as well as the positive or negative effect of each feature on patient prognosis. Mean decrease in Gini index was calculated for each feature of the RF based on the underlying decision trees [49]. Finally, since the selected SVM had a linear kernel, feature ranking coefficients were readily available [50]. A threshold was set to filter out features with low feature importance. In particular, the threshold was set to two times the mean importance of all features for the DT, LR, and RF, whereas two times the median importance was used for the LSVM. There were 8, 10, 25, and 16 important features for DT, LR, RF, LSVM, respectively (the complete feature lists for LR, LSVM, and RF are provided as Supplementary Information). A visualization of the number of intersecting features between the various submodels of our ensemble model is shown in Supplementary Figure S6.

For the LR and LSVM submodels, high density of TBs in both the invasive frontin and tumour core is highlighted as an indicator of bad prognosis. On the contrary, high density of CD8^+^, CD3^+^ and CD68^+^ cells is consistently identified as a marker of good prognosis. In addition, high number of CD3^+^, CD68^+^PD-L1^+^ and CD163^+^PD-L1^+^ cells in the invasive front as well as the presence of CD3^+^ cells within a distance of 20*µm* from TBs are associated with good prognosis. For the DT submodel, low density of CD68^+^, high PD-L1^+^ expression, and high number of TBs (all in frontout) lead to bad prognosis, whereas, given a low density of CD68^+^ and PD-L1^+^ expression in frontout, prognosis depends on the number of CD68^+^ in frontin. Finally, the majority of the patients with good prognosis had high CD68^+^ in frontout, nonzero PD-L1^+^ expression in core, low CD163^+^ in frontout, and high CD3^+^ in frontout. Similar to the previous submodels, TBs and CD68^+^ cells were the most important predictors of 5 year prognosis for RF. In addition, RF employed more spatial features than any of the other submodels including but not limited to PD-L1^+^ expression within a distance of 20*µm* from TBs and 150*µm* from M2 macrophages as well as the presence of CD3^+^ and CD8^+^ cells within a distance range of 20 − 50*µm* from TB. All of the important features are listed in Supplementary Tables S1 and S2.

## Discussion

In the last decade, advances in the rapidly growing field of tumour immunology have provided further insights into the dynamic nature of the multifaceted immune response throughout the various stages of cancer initiation, evasion, and progression. Concomitantly, multiple research groups have successfully leveraged this new knowledge to improve cancer prognosis, thereby providing evidence for the clinical relevance of immuno-oncology [12]. Accurate patient prognosis is crucial for improving the survival rates of cancer patients since it is a prerequisite to delivering the most effective treatment for each patient. In fact, multiple papers have shown that the quantitative characterization of the tumour-immune microenvironment components, including TILs, TAMs, and immune checkpoints, can yield information of prognostic relevance [51, 52, 53]. Particularly, tumour cells surrounded by a large number of prominent intra-tumoural and peri-tumoural TILs and M1 macrophages have been related to better prognosis in several types of cancer [54, 55], whereas a high content of M2 macrophages and TBs has been associated with poorer outcome [52, 56]. In addition, related research has reported the significance of PD-L1 expression on tumour tissues as an independent poor prognostic factor [57, 58]. In this paper, we have investigated the clinical relevance of immune system biomarkers, TILs, TAMs, TBs, and PD-L1, in MIBC patient prognosis.

Haematoxylin and Eosin (H&E) is still the most important and commonly used histochemical staining method for studying and diagnosing tissue diseases in histopathology. However, imaging of H&E stained FFPE tissue has limitations including the inability to quantify the complex cellular states as well as to identify distinct cell populations in the tumour-immune microenvironment. With the advent of whole slide imaging and the increasing adoption of digital pathology in the clinic [59], multiplex methodologies have the potential to provide significantly more information about the underlying tumour-immune microenvironment than single-marker (i.e. single label immunohistochemistry) and conventional histochemical staining based methodologies [60]. The development of single protein-based biomarkers to explain patient-level behaviour is hindered by the vast signalling network mediating the heterotypic cell-cell crosstalk between cancer, stromal and immune cells. Instead, with multiplexed methodologies, various proteins can be simultaneously captured on a single tissue sample, encapsulating the tumour-immune architecture from the cellular level down to the subcellular, ultimately providing more information about the microenvironment.

In our approach, multiplexed immunofluorescence was used to visualize TBs, general and cytotoxic T-cells, M1, M2, and total macrophages, and their co-expression of immune checkpoint ligand PD-L1 in order to quantify their numbers and densities, as well as their pairwise spatial distributions across defined areas (tumour core, invasive frontin and frontout) within a WSI. Over the last decade, multiple studies have investigated the topographical distribution of the immune cells within the tumour microenvironment [61, 62, 63, 64]. It is known that tumour-infiltrating immune cells are scattered in the tumour core and the invasive front whereas their density in each tumour region is correlated with patient outcome [62]. Furthermore, the analysis of multiple tumour regions (tumour core and invasive front) was shown to improve the prediction accuracy of patient survival compared to single-region analysis [11, 62]. In addition, Immunoscore, a classification system based on the quantification of two lymphocyte populations (CD3 and CD8) within the tumour core and the invasive front of tumour, has been shown to have a prognostic significance superior to that of the TNM staging system in patients with colorectal carcinoma [65, 11]. The image, spatial, and clinical features contain a multitude of information about the state of the disease and collectively portray a more holistic view of each patient pathophysiology. We hypothesized that these features can predict the aggressiveness of MIBC, and therefore suggest whether a patient should be considered low or high risk of disease-specific death.

ML contains a plethora of classifiers that have been employed with success in multiple instances, including diagnosis, segmentation, prognosis, and even therapy planning [41]. In our methodology, survival analysis is turned into a binary classification problem, thus enabling traditional ML algorithms and workflows to be readily employable. In addition, to counter the possibility of overfitting due to having a small data set yet high dimensional feature space, nested cross validation with a separate testing set was adopted. The proposed ensemble model significantly surpasses under all metrics – AUROC, Accuracy, Specificity, F1 score, Hazard ratio – (89.3%/80.0%/71.4%/83.3%/32.5) the gold standard, TNM staging (64.3%/50.0%/28.6%/44.0%/3.3), as summarized in Table 2. It consists of a LSVM that uses image features, a DT that uses image and clinical features, a LR that uses image and spatial features, and a RF that uses all features.

The results of our study suggest that the characterization of a broad immune cell population, as well as their spatial organization in relation to cancer cells, enables a better estimation of survival compared to TNM staging system in MIBC patients and provides further biological insights. In particular, we found that high content of TBs in the invasive frontin, frontout, and tumour core as well as low number of CD68^+^ cells and high PD-L1 expression in the invasive frontout are indicators of bad prognosis. This supports previous findings in the literature [33, 36, 66, 53]. High density of CD8^+^, CD3^+^, and CD68^+^ cells in the invasive frontin, frontout, and tumour core was associated with good prognosis by our models, as similarly reported in other studies [23, 67]. In addition, high number of CD3^+^ and CD68^+^ cells as well as high number of CD3^+^ cells clustered within a distance of 20*µm* from TBs were linked to good prognosis, similar to previous findings in the literature [37]. Finally, we found that high density of CD163^+^ cells without PD-L1 expression in frontout is associated with bad prognosis by the DT submodel, whereas the LSVM submodel employed high number of CD163^+^PD-L1^+^ cells in frontout as an indication of good prognosis.

In summary, we have demonstrated that ML classifiers using image and spatial features from WSIs combined with clinical features from medical records can separate MIBC patients into low and high risk groups based on a 5 year prognosis. The present approach outperforms the current clinical staging system, TNM, utilized by pathologists to determine patient prognosis. Our findings show that investigating features from the tumour-immune microenviroment in relation to survival can provide further insights for histopathological studies, thereby contributing to better ways for predicting survivability and enabling better quality of care.

## Methods

### Patients and tissue samples

Tissue specimens from patients with MIBC who underwent radical cystectomy at Royal Infirmary and Western General Hospital in Edinburgh between the years 2006 to 2013 were collated into a cohort. Patients were excluded from this study either due to incomplete clinical records, extensive tissue section artefacts, or data censoring. The final study cohort was comprised of 78 patients. Archived FFPE tissue blocks presenting the deepest invasion of cancer were selected for each patient based on the assessment of haematoxylin and eosin (H&E) stained slides by a pathologist and a research scientist. The corresponding unstained tissue sections were collected from the NHS Lothian NRS BioResource Research Tissue Bank (Ethical status/approval ref: 10/S1402/33), conforming to protocols approved by the East of Scotland Research Ethics Service. Clinicopathological data that included age, sex and TNM stage status along with survival data was retrieved from available electronic medical records. Patients were followed up for a total time of 113 months with a median survival time of 23.6 months. In order to maintain the anonymity of the patient information, the samples had been de-identified prior to conducting this study.

### Multiplex immunofluorescence and whole slide imaging

For each patient, automated tyramide-based immunofluorescence was performed on two de-paraffinised serial 3*µm* thick sections of FFPE tissue mounted on superfrost plus slides using a Dako link 48 autostainer (Dako, Agilent Technologies). Primary antibodies against Pan-cytokeratin (PanCK), CD3, CD8, CD68, CD163 and PD-L1 were used to label urothelial cells, general T-cells, cytotoxic T-cells, M1/M2 (total) macrophages, M2 macrophages and immune checkpoint ligand PD-L1, respectively (see Supplementary Table S5 for further details about the primary antibodies). To increase the detection sensitivity and to visualise the target protein, TSA reagents FITC, CY3 and CY5 were used for CD3, CD8, and PD-L1 respectively. Alexa Fluor 750 conjugated streptavidin antibody was used for the detection of PanCK. See Supplementary Table S6 for the detection and visualization reagents. Nuclei were counterstained with Hoechst (Hoechst 33342, Cat# H3570, ThermoFisher Scientific) and ProLong Gold Antifade mountant (Cat# P36930, ThermoFisher Scientific) applied directly to the tissue samples. The multiplex immuno-labeled tissue slides were scanned at 20× magnification and digitized into whole slide fluorescence images using a Carl Zeiss AxioScan.Z1 scanner (Zeiss, Göttingen, Germany). Examples are shown in Figure 2. Supplementary Table S7 and Supplementary Figure S7 show the excitation and emission wavelengths along with the exposure time for each antibody used for multispectral imaging.

**Figure 2:**
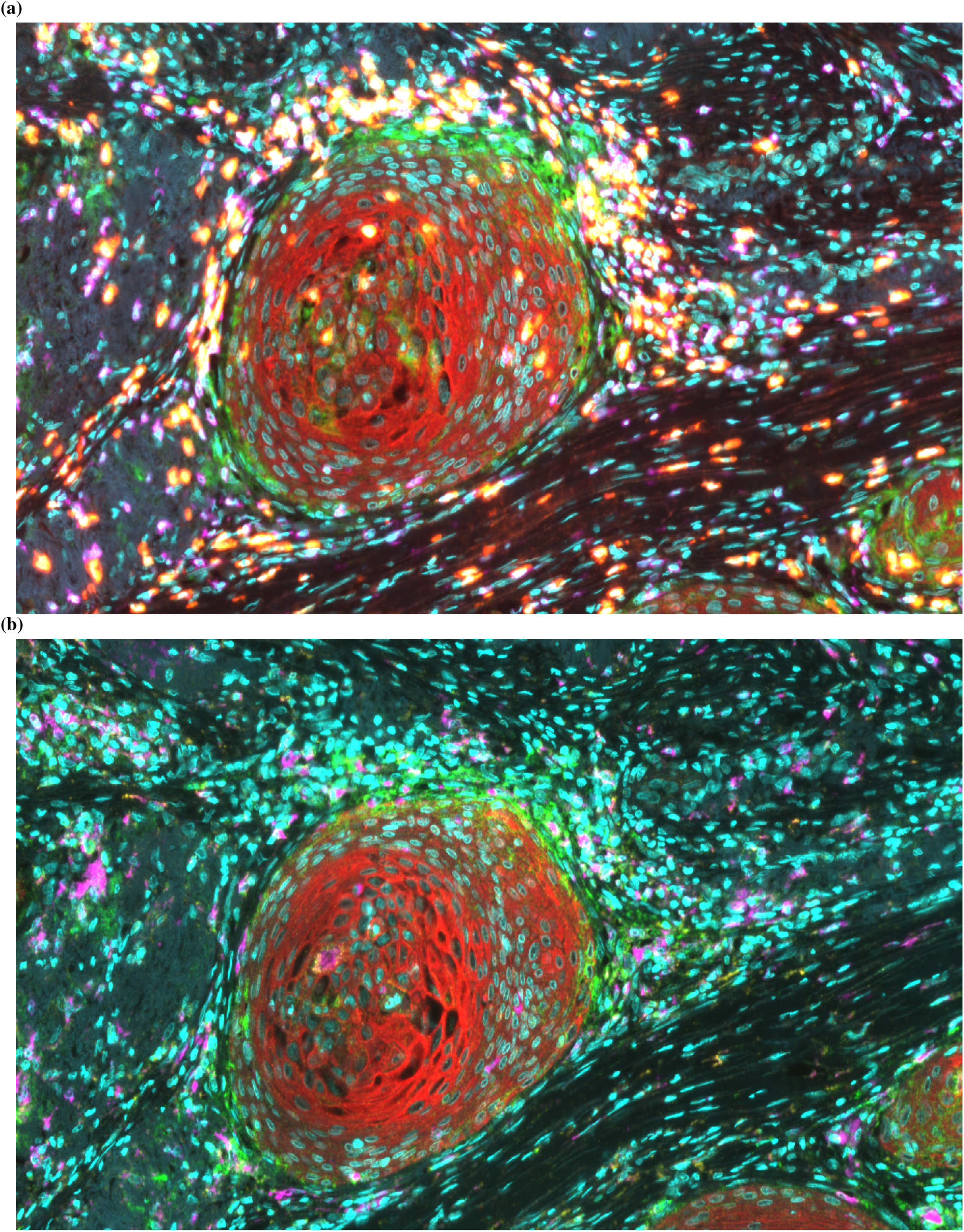
Examples of (a) TILs (*Nuclei: Cyan, Cancer: Red , PD-L1: Green, CD3: Purple, CD8: Yellow*) and (b) TAMs (*Nuclei: Cyan, Cancer: Red , PD-L1: Green, CD68: Purple, CD163: Yellow*) visualized using multiplexed immunofluorescence.

### Detection of cell nuclei

For nucleus detection, the methodology described by Brieu and Schmidt [68] was adopted. For completeness, a high level description of this methodology is provided. The methodology is comprised of four distinct stages: (a) A classification RF was trained with long-range spatial context features [69] extracted from manually annotated epithelial and non-epithelial regions (*n* = 4, resolution=1200 × 1200 pixels). Subsequently, the RF generated an epithelial mask for all IF images [70]. (b) A regression RF was trained to generate proximity maps using coordinates from manual annotations of cell nuclei (*n* = 750 from 19 fields of view, resolution=1485 × 1485). Intuitively, a proximity map encloses the distance to the closest nucleus center for each pixel of the input image. (c) A regression RF was trained to generate surface area maps using manual annotations (*n* = 250 from 8 fields of view, resolution=580 × 580 pixels) [68]. A surface area model provides a mask of the initial image wherein each pixel is either zero, if it is not a part of a nucleus, or a positive real number, if it is part of a nucleus. The positive real number is the area of the corresponding nucleus. (d) At this stage, nuclei centers (see Supplementary Figure S3) are localized based on the proximity and surface area maps that were generated in (b) and (c) [68]. Note that both the original image and the corresponding epithelial mask (produced in (a)) are given as inputs to the models in (b) and (c) using only the PanCK and Hoechst IF channels. Data augmentation with varying scale, rotation, as well as intensity for both PanCK and Hoechst IF channels was implemented in all stages. Finally, regions with necrotic tissue and any type of artefact, such as autofluorescence, were not included in any of the training data.

### Segmentation of epithelial cells for the identification of tumour buds

For the quantification of tumours buds, segmentation of the epithelial cells was required. The CNN-RF methodology described by Brieu et al [71] and extended in [36] was adopted. Briefly, a convolutional neural network (CNN) was trained on an annotated data set of epithelium and non-epithelium IF images. The data set comprised of 142 × 142 pixel regions that were selected by an expert. The Hoechst, PanCK, CD3 and CD8 IF channels were normalised following the approach described by Brieu et al. [71]. An intensity interval was imposed for the PD-L1, CD163, and CD68 IF channels by computing minimum and maximum values from the segmented epithelium regions of each slide. Once trained, the CNN produced a coarse segmentation mask of epithelium regions. Normalized PanCK, Hoechst, CD3, and CD8 channels of the IF images were used as input to the CNN. The predicted epithelium probability layer was used together with the original IF channels as input to a RF. Finer-grained segmentation masks were produced by the RF, enabling a more accurate segmentation of the epithelium. As detailed in previous work [36], the output of the CNN-RF is finally ensembled with the output of a semantic segmentation network [72] to generate the final epithelium segmentation results. Tumour buds were classified as epithelium objects containing one to four nuclei [36]. A total number of 97262 patches were used for training and 9742 patches for validation of the CNN-RF model. The semantic segmentation network was trained with 19093 patches and validated with 6987 patches with 256 × 256 pixels.

### Cell classification

For cell classification, given the normalised IF channels, a circular neighbourhood of each cell nuclei is defined (11 × 11 pixel radius) and the mean normalised intensity of the neighbourhood is computed for each IF marker (CD3, CD8, CD68, CD163, PD-L1 and PanCK). Cells are classified as positive or negative for a given IF marker if the corresponding mean normalized intensity is above or below a determined threshold, respectively. In our experiments, the threshold for all the IF markers was set to 32/256 = 0.125.

### Pairwise spatial distributions of lymphocytes, macrophages, tumour buds and PD-L1

The extracted point coordinates of cell nuclei and immune checkpoint ligand PD-L1 expression were localized across the WSIs as shown in Supplementary Figure S1. The Ripley’s K function [73] was adopted for investigating how TBs, PD-L1, and the different populations of immune cells are distributed around each other. In particular, given two populations *X* and *Y* , Ripley’s K function estimates the density of *Y* within a circle of radius *r* around points *X*. As illustrated in Figure 3, assuming a Poisson distribution, the Ripley’s K function can identify whether a population *Y* is dispersed, randomly distributed, or clustered around another population *X*. The *K* function is given as:

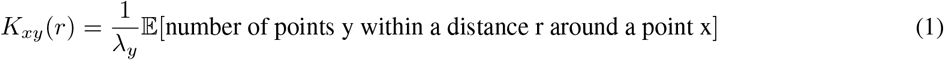

where 𝔼[·] encloses all of the points of type y within a distance r of a randomly selected point of type x. Theoretically, if the point pattern of points *Y* around *X* follows complete spatial randomness, also known as a homogeneous Poisson process, the value of K function is *πr*^2^. The L function [74] is a modification of equation 1, so that the expected output value is *r*, i.e.:

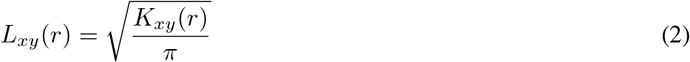

**Figure 3:**
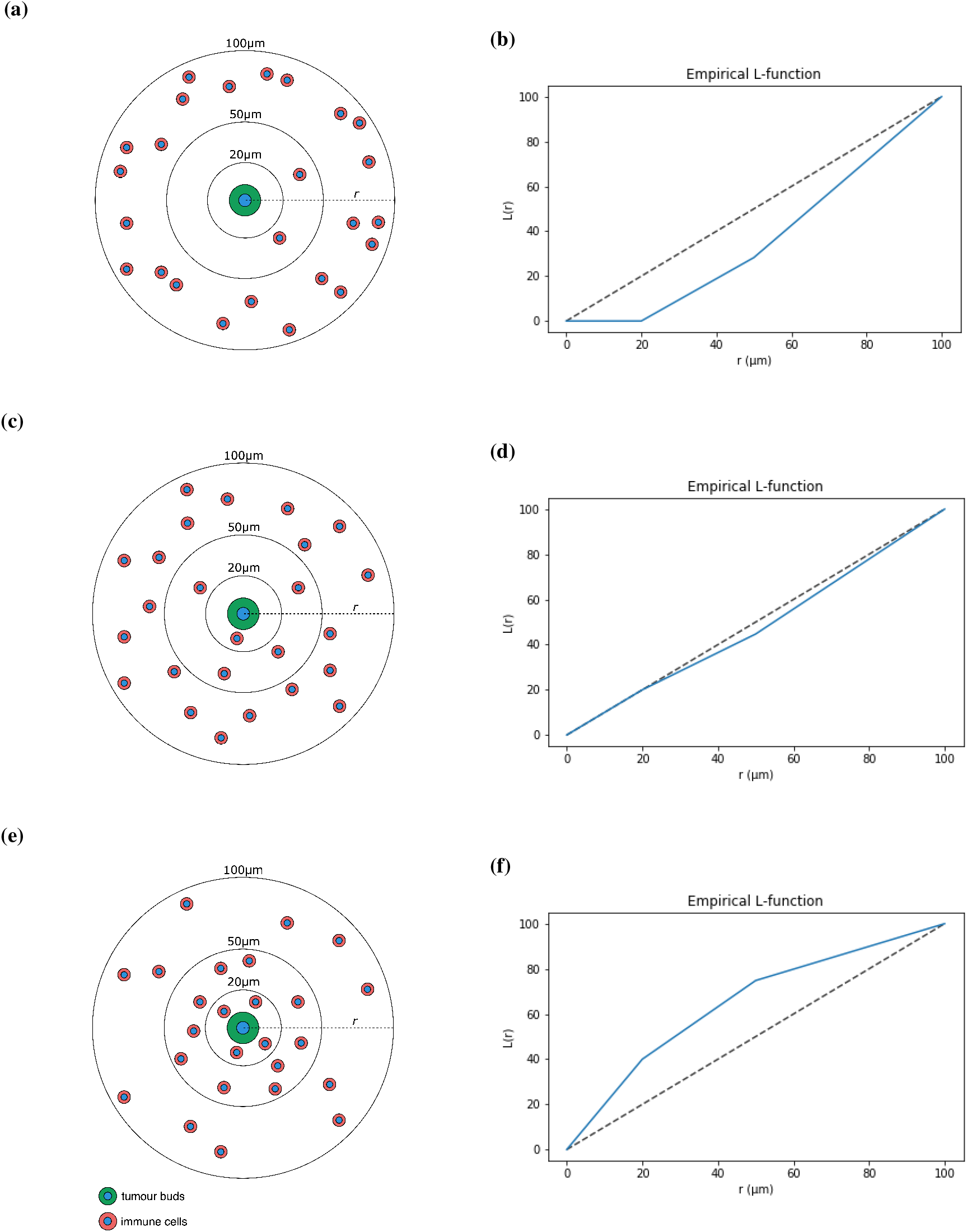
Schematic representation of different immune cell distributions from the nuclear centre of a tumour bud (a, c, e), and their corresponding L function values at different radii (b, d, f). The immune cell population is either (a–b) dispersed, (c–d) randomly distributed, or (d–e) clustered around the TB.

This enables a more intuitive interpretation of the function’s output value in relation to *r*. The *L* function was calculated for TAMs, TILs, and PD-L1 surrounding TBs as well as PD-L1 surrounding TAMs and TILs for a series of increasing distances *r* where *r* ∈ {20, 50, 100, 150, 200, 250} *µ*m. While some approaches calculate the area under the curve of the *L* function against different *r* values [75], we provided the pairwise spatial distributions between PD-L1, TBs, and the immune populations directly to the ML classifiers as distinct features.

### Binary survival analysis

Survival analysis is broadly defined as the analysis of data that involve the time to the occurrence of an event of interest [76]. Herein, the event of interest was the death of an individual due to MIBC. A characteristic of survival analysis is censoring. In our cohort, some patients were right-censored either because the end of study was reached and the event of interest did not occur, or because the patients succumbed to a cause other than MIBC (abbreviated as OTD-censoring) [76].

Patient survivability was binarized based on a specific time cutoff. Similarly to previous work, a 5 year prognosis was investigated [42, 77, 78]. Patients that succumbed to MIBC within 5 years were denoted as patients with a bad prognosis whereas those that survived the 5 year cutoff were denoted as patients with a good prognosis. Inevitably, patients that died to an unrelated cause prior to the prognostic cutoff, i.e. they were part of the OTD-censored data, had to be excluded (19% patient exclusion). It is worth mentioning that removing these patients does not introduce bias since time to censoring was random, i.e. OTD-censoring was not known *a priori*. A consequence of this approach is that survival analysis was turned into a binary classification problem. Furthermore, due to removing censoring, traditional ML models were readily employable.

### Model selection, algorithm selection, and performance evaluation

Both model selection and algorithm selection attempt to collectively maximize the predictive performance of the final ML model. However, ML algorithms are prone to overfitting, i.e. in finding and using patterns which arise from noise in the data. Such noisy patterns do not generally extend beyond the specific data set since noise is typically random. With both a small data set and a complicated model, the likelihood of overfitting increases. Testing the performance of a trained ML model on unseen data constitutes the mainstay in evaluating the generalizability of a ML model, and therefore, in identifying whether a model has overfitted. As such, a subset of the initial cohort was kept aside as the testing data set. In particular, using stratified random sampling, two subsets were created, the training set with 75% of the initial data (58 patients), and the testing set with 25% (20 patients). The testing set was only used at the performance evaluation stage to avoid introducing bias to the generalization performance estimate.

Traditionally, model selection is the process in which a ML algorithm is configured. Most ML algorithms come with a number of configuration variables, commonly referred to as hyperparameters. Even though common hyperparameter configurations can be employed, it has been observed that hyperparameter tuning for a specific task can be the key between chance and state-of-the-art models [79]. Since manual tuning can be time consuming and counter-intuitive in high dimensional spaces, most ML methodologies adopt automated hyperparameter tuning.

One of the most popular approaches to model selection constitutes grid search where each hyperparameter is given a predefined list of values, and the best hyperparameter configuration is selected after evaluating all the combinations. For example, given hyperparameters *A* and *B*, and lists *V*_*A*_ = [1, 10, 100] and *V*_*B*_ = [0.1, 0.5], the following combinations were evaluated under grid search: ∀(*A, B*) ∈ [(1, 0.1), (1, 0.5), (10, 0.1), (10, 0.5), (100, 0.1), (100, 0.5)]. However, as shown by Bergstra and Bengio [80], random sampling provides a better tuning strategy.

In random search, a number of hyperparameter configurations are evaluated by sampling from predefined hyperparameter distributions and densities. For example, given hyperparameters *A* and *B* with *D*_*A*_ ~ *N* (0, 1) and *V*_*B*_ = [0.1, 0.5], where *D*_*A*_ is a standard Gaussian distribution, the following five combinations could have been sampled and evaluated under random search: ∀(*A, B*) ∈ [(−0.12, 0.1), (−0.14, 0.5), (−0.94, 0.5), (0.44, 0.1), (−1.3, 0.5)]. In our methodology, 200 hyperparameter configurations were randomly sampled and evaluated for each ML algorithm. A table of the distributions and densities used is provided in Supplementary Information S4.

Not every ML algorithm will perform equally well on different problems and different data. In addition, there is no theoretical ranking suggesting that one algorithm is better than another [81]. Hence, similar to hyperparameter tuning, algorithm selection is yet another meta-optimization task that needs to be performed for maximizing predictive performance. However, as argued by Bergstra et al. [79]: “Since the performance of a given technique depends on both the fundamental quality of the algorithm and the details of its tuning, it is sometimes difficult to know whether a given technique is genuinely better, or simply better tuned.” Consequently, algorithm selection should involve model selection. Therefore, each ML algorithm was first tuned using fivefold cross validation and then compared against each other using twofold cross validation. This nested cross validation translates to optimizing the hyperparameters of each ML algorithm twice, and then measuring their performance on the corresponding evaluation folds (see Figure 4). Subsequently, the ML algorithm which performed better than the rest across two different training and validation folds is selected. It is important to highlight how in most cases each ML algorithm is evaluated based on two different hyperparameter configurations. Nevertheless, once the ML algorithm has been selected, yet another hyperparameter tuning phase is implemented to find an optimal hyperparameter configuration based on the whole training data set. Five ML algorithms with different theoretical underpinnings were selected to compete against each other; Decision Tree (DT), Random Forest (RF), Support Vector Machine (SVM), Logistic Regression (LR), and *k* nearest neighbours (KNN). Finally, a preprocessing step of feature normalisation was added to all classifiers except DT and RF.

**Figure 4:**
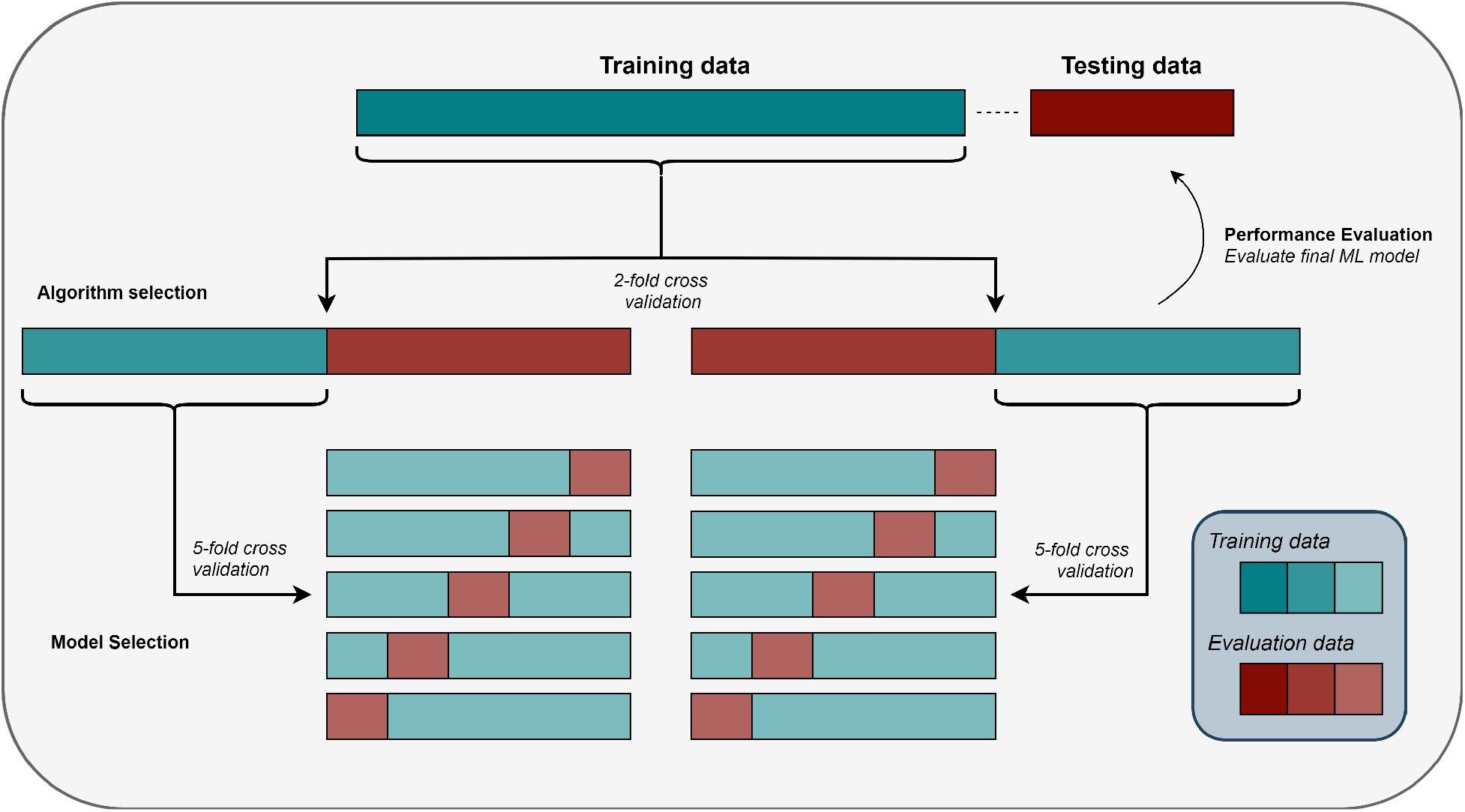
Pictorial representation of nested cross validation with an independent testing set. (A) Performance Evaluation: The best ML algorithm (selected by the outer cross validation, see *B*) was trained on the training data set and subsequently evaluated on the testing data set. (B) Algorithm Selection: Each ML algorithm (with hyperparameters tuned based on the inner cross validation, see *C*) was trained and tested on the corresponding training and evaluation folds respectively. The best ML algorithm was selected based on the average performance of both evaluation folds. (C) Model Selection: ML models with randomly sampled hyperparameter configurations were trained and tested based on a fivefold cross validation. The best hyperparameter configuration for each ML algorithm was selected based on the performance on all five evaluation folds.

### Stratified sampling

In order to avoid sampling subsets with different class distributions (classes are based on survival with a 5 year cutoff) than the original cohort, stratified sampling was used. Intuitively, when sampling from a data set with stratification, proportionally many patients from each class are sampled. Given a data set with 75 patients of class *C*_1_ and 25 patients of class *C*_2_, a 20% sample would contain 15 patients from class *C*_1_ and 5 patients from class *C*_2_.

## Supporting information

Feature list

Logistic regression feature importance

Random forest feature importance

Support vector machine feature importance

## Data availability

The data and the code of this study are available from the corresponding authors upon request.

## Acknowledgements

We would like to thank Ms Frances Rae, Dr Marie O’Donnell, and the NHS Lothian Tissue Governance Unit for providing the patient clinical data and the tissue samples.

## Author information

### Contributions

CGG, PDC and GS conceived the idea of this project. DJH and PDC selected the patient samples. CGG and IPN carried out the web-lab experiments. CGG, ND, NB, PDC designed different parts of the methodology. CGG, ND and NB implemented the methodology and conducted the corresponding experiments. The manuscript was written by CGG and ND, and revised by all authors. DJH provided pathology expertise. OA, GS, and PDC supervised this project.

## References

[1] Oner Sanli, Jakub Dobruch, Margaret A Knowles, Maximilian Burger, Mehrdad Alemozaffar, Matthew E Nielsen, and Yair Lotan. Bladder cancer. Nature reviews Disease primers, 3:17022, 2017.

[2] Ashish M Kamat, Noah M Hahn, Jason A Efstathiou, Seth P Lerner, Per-Uno Malmström, Woonyoung Choi, Charles C Guo, Yair Lotan, and Wassim Kassouf. Bladder cancer. The Lancet, 388(10061):2796–2810, 2016.

[3] Margaret A Knowles and Carolyn D Hurst. Molecular biology of bladder cancer: new insights into pathogenesis and clinical diversity. Nature Reviews Cancer, 15(1):25–41, 2015.

[4] Maria Frantzi, Kim E Van Kessel, Ellen C Zwarthoff, Mirari Marquez, Marta Rava, Núria Malats, Axel S Merseburger, Ioannis Katafigiotis, Konstantinos Stravodimos, William Mullen, et al. Development and validation of urine-based peptide biomarker panels for detecting bladder cancer in a multi-center study. Clinical Cancer Research, 22(16):4077–4086, 2016.

[5] Sandip M Prasad, G Joel DeCastro, and Gary D Steinberg. Urothelial carcinoma of the bladder: definition, treatment and future efforts. Nature Reviews Urology, 8(11):631, 2011.

[6] A Gordon Robertson, Jaegil Kim, Hikmat Al-Ahmadie, Joaquim Bellmunt, Guangwu Guo, Andrew D Cherniack, Toshinori Hinoue, Peter W Laird, Katherine A Hoadley, Rehan Akbani, et al. Comprehensive molecular characterization of muscle-invasive bladder cancer. Cell, 171(3):540–556, 2017.

[7] Alexander P Glaser, Damiano Fantini, Ali Shilatifard, Edward M Schaeffer, and Joshua J Meeks. The evolving genomic landscape of urothelial carcinoma. Nature Reviews Urology, 14(4):215, 2017.

[8] Eva Compérat, Justine Varinot, Julien Moroch, Caroline Eymerit-Morin, and Fadi Brimo. A practical guide to bladder cancer pathology. Nature Reviews Urology, 15(3):143, 2018.

[9] American Joint Committee on Cancer. AJCC - Cancer Staging Manual. https://cancerstaging.org/references-tools/deskreferences/Pages/default.aspx.

[10] Constantine Alifrangis, Ursula McGovern, Alex Freeman, Thomas Powles, and Mark Linch. Molecular and histopathology directed therapy for advanced bladder cancer. Nature Reviews Urology, 16(8):465–483, 2019.

[11] Jérôme Galon, Bernhard Mlecnik, Gabriela Bindea, Helen K Angell, Anne Berger, Christine Lagorce, Alessandro Lugli, Inti Zlobec, Arndt Hartmann, Carlo Bifulco, et al. Towards the introduction of the ‘immunoscore’in the classification of malignant tumours. The Journal of pathology, 232(2):199–209, 2014.

[12] Franck Pagès, Bernhard Mlecnik, Florence Marliot, Gabriela Bindea, Fang-Shu Ou, Carlo Bifulco, Alessandro Lugli, Inti Zlobec, Tilman T Rau, Martin D Berger, et al. International validation of the consensus immunoscore for the classification of colon cancer: a prognostic and accuracy study. The Lancet, 391(10135):2128–2139, 2018.

[13] Tom Donnem, TK Kilvaer, S Andersen, E Richardsen, EE Paulsen, SM Hald, S Al-Saad, Odd Terje Brustugun, A Helland, M Lund-Iversen, et al. Strategies for clinical implementation of tnm-immunoscore in resected nonsmall-cell lung cancer. Annals of Oncology, 27(2):225–232, 2015.

[14] Annacarmen Petrizzo and Luigi Buonaguro. Application of the immunoscore as prognostic tool for hepatocellular carcinoma. Journal for immunotherapy of cancer, 4(1):71, 2016.

[15] Frances R Balkwill, Melania Capasso, and Thorsten Hagemann. The tumor microenvironment at a glance. J Cell Sci, 125:5591–5596, 2012.

[16] Melissa R Junttila and Frederic J de Sauvage. Influence of tumour micro-environment heterogeneity on therapeutic response. Nature, 501(7467):346, 2013.

[17] Shelly Maman and Isaac P Witz. A history of exploring cancer in context. Nature Reviews Cancer, 18(6):359, 2018.

[18] Seok-Hyun Kim, Se-Il Go, Dae Hyun Song, Sung Woo Park, Hye Ree Kim, Inseok Jang, Jong Duk Kim, Jong Sil Lee, and Gyeong-Won Lee. Prognostic impact of cd8 and programmed death-ligand 1 expression in patients with resectable non-small cell lung cancer. British journal of cancer, 120(5):547, 2019.

[19] Friedrich Foerster, Moritz Hess, Aslihan Gerhold-Ay, Jens Uwe Marquardt, Diana Becker, Peter Robert Galle, Detlef Schuppan, Harald Binder, and Ernesto Bockamp. The immune contexture of hepatocellular carcinoma predicts clinical outcome. Scientific reports, 8(1):5351, 2018.

[20] Max D Wellenstein and Karin E de Visser. Cancer-cell-intrinsic mechanisms shaping the tumor immune landscape. Immunity, 48(3):399–416, 2018.

[21] Wolf H Fridman, Laurence Zitvogel, Catherine Sautès-Fridman, and Guido Kroemer. The immune contexture in cancer prognosis and treatment. Nature reviews Clinical oncology, 14(12):717, 2017.

[22] Y Ino, R Yamazaki-Itoh, K Shimada, M Iwasaki, T Kosuge, Yae Kanai, and N Hiraoka. Immune cell infiltration as an indicator of the immune microenvironment of pancreatic cancer. British journal of cancer, 108(4):914, 2013.

[23] Tristan A Barnes and Eitan Amir. Hype or hope: the prognostic value of infiltrating immune cells in cancer. British journal of cancer, 117(4):451, 2017.

[24] Philipp Lohneis, Marianne Sinn, Fritz Klein, Sven Bischoff, Jana K Striefler, Lilianna Wislocka, Bruno V Sinn, Uwe Pelzer, Helmut Oettle, Hanno Riess, et al. Tumour buds determine prognosis in resected pancreatic ductal adenocarcinoma. British journal of cancer, 118(11):1485, 2018.

[25] Ling Li, Ruifang Sun, Yi Miao, Thai Tran, Lisa Adams, Nathan Roscoe, Bing Xu, Ganiraju C Manyam, Xiaohong Tan, Hongwei Zhang, et al. Pd-1/pd-l1 expression and interaction by automated quantitative immunofluorescent analysis show adverse prognostic impact in patients with diffuse large b-cell lymphoma having t-cell infiltration: a study from the international dlbcl consortium program. Modern Pathology, page 1, 2019.

[26] Jean-David Fumet, Corentin Richard, Fanny Ledys, Quentin Klopfenstein, Philippe Joubert, Bertrand Routy, Caroline Truntzer, Andréanne Gagné, Marc-André Hamel, Camila Figueiredo Guimaraes, et al. Prognostic and predictive role of cd8 and pd-l1 determination in lung tumor tissue of patients under anti-pd-1 therapy. British journal of cancer, 119(8):950, 2018.

[27] Ellen L Goode, Matthew S Block, Kimberly R Kalli, Robert A Vierkant, Wenqian Chen, Zachary C Fogarty, Aleksandra Gentry-Maharaj, Aleksandra Tołoczko, Alexander Hein, Aliecia L Bouligny, et al. Dose-response association of cd8+ tumor-infiltrating lymphocytes and survival time in high-grade serous ovarian cancer. JAMA oncology, 3(12):e173290–e173290, 2017.

[28] Luca Cassetta and Jeffrey W Pollard. Targeting macrophages: therapeutic approaches in cancer. Nature Reviews Drug Discovery, 2018.

[29] David G DeNardo and Brian Ruffell. Macrophages as regulators of tumour immunity and immunotherapy. Nature Reviews Immunology, 19(6):369–382, 2019.

[30] Subhra K Biswas and Alberto Mantovani. Macrophage plasticity and interaction with lymphocyte subsets: cancer as a paradigm. Nature immunology, 11(10):889, 2010.

[31] Inti Zlobec and Alessandro Lugli. Tumour budding in colorectal cancer: molecular rationale for clinical translation. Nat Rev Cancer, 18:203–4, 2018.

[32] H Ueno, J Murphy, JR Jass, H Mochizuki, and IC Talbot. Tumourbudding’as an index to estimate the potential of aggressiveness in rectal cancer. Histopathology, 40(2):127–132, 2002.

[33] FJA Gujam, DC McMillan, ZMA Mohammed, J Edwards, and JJ Going. The relationship between tumour budding, the tumour microenvironment and survival in patients with invasive ductal breast cancer. British journal of cancer, 113(7):1066, 2015.

[34] Linde De Smedt, Sofie Palmans, Daan Andel, Olivier Govaere, Bram Boeckx, Dominiek Smeets, Eva Galle, Jasper Wouters, David Barras, Madeleine Suffiotti, et al. Expression profiling of budding cells in colorectal cancer reveals an emt-like phenotype and molecular subtype switching. British journal of cancer, 116(1):58, 2017.

[35] Alessandro Lugli, Richard Kirsch, Yoichi Ajioka, Fred Bosman, Gieri Cathomas, Heather Dawson, Hala El Zimaity, Jean-François Fléjou, Tine Plato Hansen, Arndt Hartmann, et al. Recommendations for reporting tumor budding in colorectal cancer based on the international tumor budding consensus conference (itbcc) 2016. Modern pathology, 30(9):1299, 2017.

[36] Nicolas Brieu, Christos G Gavriel, Ines P Nearchou, David J Harrison, Günter Schmidt, and Peter D Caie. Automated tumour budding quantification by machine learning augments tnm staging in muscle-invasive bladder cancer prognosis. Scientific reports, 9(1):5174, 2019.

[37] Ines P Nearchou, Kate Lillard, Christos G Gavriel, Hideki Ueno, David J Harrison, and Peter D Caie. Automated analysis of lymphocytic infiltration, tumor budding, and their spatial relationship improves prognostic accuracy in colorectal cancer. Cancer immunology research, 7(4):609–620, 2019.

[38] Drew M Pardoll. The blockade of immune checkpoints in cancer immunotherapy. Nature Reviews Cancer, 12(4):252, 2012.

[39] Chong Sun, Riccardo Mezzadra, and Ton N Schumacher. Regulation and function of the pd-l1 checkpoint. Immunity, 48(3):434–452, 2018.

[40] KH Yu, C Zhang, G. J. Berry, R. B. Altman, C. Ré, D. L. Rubin, and M. Snyder. Predicting non-small cell lung cancer prognosis by fully automated microscopic pathology image features. Nature Communications, 7(1), 2016.

[41] A. Madabhushi and G. Lee. Image analysis and machine learning in digital pathology: Challenges and opportunities. Medical Image Analysis, 33:170–175, 2016.

[42] N. Dimitriou, O. Arandjelovicć, D. J. Harrison, and P. D. Caie. A principled machine learning framework improves accuracy of stage II colorectal cancer prognosis. npj Digital Medicine, 1(1), 2018.

[43] R. Mokarram and M. Emadi. Classification in non-linear survival models using cox regression and decision tree. Annals of Data Science, 4(3):329–340, 2017.

[44] P. Wang, Y. Li, and C. K. Reddy. Machine learning for survival analysis: A survey, 2017.

[45] Nathalie Harder, Maria Athelogou, Harald Hessel, Nicolas Brieu, Mehmet Yigitsoy, Johannes Zimmermann, Martin Baatz, Alexander Buchner, Christian G Stief, Thomas Kirchner, et al. Tissue phenomics for prognostic biomarker discovery in low-and intermediate-risk prostate cancer. Scientific reports, 8(1):4470, 2018.

[46] Gerd Binnig, Ralf Huss, and Günter Schmidt. Tissue phenomics: Profiling cancer patients for treatment decisions. CRC Press, 2018.

[47] Maria Athelogou, Günter Schmidt, Arno Schäpe, Martin Baatz, and Gerd Binnig. Cognition network technology–a novel multimodal image analysis technique for automatic identification and quantification of biological image contents. In Imaging cellular and molecular biological functions, pages 407–422. Springer, 2007.

[48] S. Raschka. Model evaluation, model selection, and algorithm selection in machine learning, 2018.

[49] G. Louppe, L. Wehenkel, A. Sutera, and P. Geurts. Understanding variable importances in forests of randomized trees. In C. J. C. Burges, L. Bottou, M. Welling, Z. Ghahramani, and K. Q. Weinberger, editors, Advances in Neural Information Processing Systems 26, pages 431–439. Curran Associates, Inc., 2013.

[50] I. Guyon, J. Weston, S. Barnhill, and V. Vapnik. Gene selection for cancer classification using support vector machines. Machine Learning, 46(1):389–422, Jan 2002.

[51] Marloes JM Gooden, Geertruida H de Bock, Ninke Leffers, Toos Daemen, and Hans W Nijman. The prognostic influence of tumour-infiltrating lymphocytes in cancer: a systematic review with meta-analysis. British journal of cancer, 105(1):93, 2011.

[52] Taisuke Yagi, Yoshifumi Baba, Kazuo Okadome, Yuki Kiyozumi, Yukiharu Hiyoshi, Takatsugu Ishimoto, Masaaki Iwatsuki, Yuji Miyamoto, Naoya Yoshida, Masayuki Watanabe, et al. Tumour-associated macrophages are associated with poor prognosis and programmed death ligand 1 expression in oesophageal cancer. European Journal of Cancer, 111:38–49, 2019.

[53] Manuel D Keller, Christina Neppl, Yasin Irmak, Sean R Hall, Ralph A Schmid, Rupert Langer, and Sabina Berezowska. Adverse prognostic value of pd-l1 expression in primary resected pulmonary squamous cell carcinomas and paired mediastinal lymph node metastases. Modern pathology, 31(1):101, 2018.

[54] Yohei Masugi, Tokiya Abe, Akihisa Ueno, Yoko Fujii-Nishimura, Hidenori Ojima, Yutaka Endo, Yusuke Fujita, Minoru Kitago, Masahiro Shinoda, Yuko Kitagawa, et al. Characterization of spatial distribution of tumorinfiltrating cd8+ t cells refines their prognostic utility for pancreatic cancer survival. Modern Pathology, page 1, 2019.

[55] CM Ohri, A Shikotra, RH Green, DA Waller, and P Bradding. Macrophages within nsclc tumour islets are predominantly of a cytotoxic m1 phenotype associated with extended survival. European Respiratory Journal, 33(1):118–126, 2009.

[56] Ailín C Rogers, David Gibbons, Ann M Hanly, John MP Hyland, P Ronan O’connell, Desmond C Winter, and Kieran Sheahan. Prognostic significance of tumor budding in rectal cancer biopsies before neoadjuvant therapy. Modern Pathology, 27(1):156, 2014.

[57] Song Xue, Ge Song, and Jinming Yu. The prognostic significance of pd-l1 expression in patients with glioma: a meta-analysis. Scientific reports, 7(1):4231, 2017.

[58] Cuiling Zhou, Jianjun Tang, Huanhuan Sun, Xiaobin Zheng, Zhanyu Li, Tiantian Sun, Jie Li, Shuncong Wang, Xiuling Zhou, Hongliu Sun, et al. Pd-l1 expression as poor prognostic factor in patients with non-squamous non-small cell lung cancer. Oncotarget, 8(35):58457, 2017.

[59] Peter Caie, N Dimitriou, IP Nearchou, O Arandjelovic, and D Harrison. Artificial intelligence driving automated pathology: icaird and beyond. In VIRCHOWS ARCHIV, volume 475, pages S60–S60, 2019.

[60] Edwin Roger Parra, Alejandro Francisco-Cruz, and Ignacio Ivan Wistuba. State-of-the-art of profiling immune contexture in the era of multiplexed staining and digital analysis to study paraffin tumor tissues. Cancers, 11(2):247, 2019.

[61] Jakob Nikolas Kather, Meggy Suarez-Carmona, Pornpimol Charoentong, Cleo-Aron Weis, Daniela Hirsch, Peter Bankhead, Marcel Horning, Dyke Ferber, Ivan Kel, Esther Herpel, et al. Topography of cancer-associated immune cells in human solid tumors. Elife, 7:e36967, 2018.

[62] Jérôme Galon, Anne Costes, Fatima Sanchez-Cabo, Amos Kirilovsky, Bernhard Mlecnik, Christine Lagorce-Pagès, Marie Tosolini, Matthieu Camus, Anne Berger, Philippe Wind, et al. Type, density, and location of immune cells within human colorectal tumors predict clinical outcome. Science, 313(5795):1960–1964, 2006.

[63] Gabriela Bindea, Bernhard Mlecnik, Marie Tosolini, Amos Kirilovsky, Maximilian Waldner, Anna C Obenauf, Helen Angell, Tessa Fredriksen, Lucie Lafontaine, Anne Berger, et al. Spatiotemporal dynamics of intratumoral immune cells reveal the immune landscape in human cancer. Immunity, 39(4):782–795, 2013.

[64] Lisa König, Fabian D Mairinger, Oliver Hoffmann, Ann-Kathrin Bittner, Kurt W Schmid, Rainer Kimmig, Sabine Kasimir-Bauer, and Agnes Bankfalvi. Dissimilar patterns of tumor-infiltrating immune cells at the invasive tumor front and tumor center are associated with response to neoadjuvant chemotherapy in primary breast cancer. BMC cancer, 19(1):120, 2019.

[65] Jérôme Galon, Franck Pagès, Francesco M Marincola, Helen K Angell, Magdalena Thurin, Alessandro Lugli, Inti Zlobec, Anne Berger, Carlo Bifulco, Gerardo Botti, et al. Cancer classification using the immunoscore: a worldwide task force. Journal of translational medicine, 10(1):205, 2012.

[66] Sari Riihijärvi, Idun Fiskvik, Minna Taskinen, Heli Vajavaara, Maria Tikkala, Olav Yri, Marja-Liisa Karjalainen-Lindsberg, Jan Delabie, Erlend Smeland, Harald Holte, et al. Prognostic influence of macrophages in patients with diffuse large b-cell lymphoma: a correlative study from a nordic phase ii trial. haematologica, 100(2):238–245, 2015.

[67] Nathalie Chaput, Magali Svrcek, Anne Aupérin, Clara Locher, Françoise Drusch, David Malka, Julien Taïeb, Diane Goéré, Michel Ducreux, and Valérie Boige. Tumour-infiltrating cd68+ and cd57+ cells predict patient outcome in stage ii–iii colorectal cancer. British journal of cancer, 109(4):1013–1022, 2013.

[68] Nicolas Brieu and Günter Schmidt. Learning size adaptive local maxima selection for robust nuclei detection in histopathology images. In 2017 IEEE 14th International Symposium on Biomedical Imaging (ISBI 2017), pages 937–941, 2017.

[69] Antonio Criminisi, Jamie Shotton, and Stefano Bucciarelli. Decision forests with long-range spatial context for organ localization in ct volumes. In Medical Image Computing and Computer-Assisted Intervention (MICCAI), pages 69–80, 2009.

[70] Nicolas Brieu, Olivier Pauly, Johannes Zimmermann, Gerd Binnig, and Günter Schmidt. Slide-specific models for segmentation of differently stained digital histopathology whole slide images. In Medical Imaging 2016: Image Processing, volume 9784, page 978410, 2016.

[71] Nicolas Brieu, Christos G Gavriel, David J Harrison, Peter D Caie, and Günter Schmidt. Context-based interpolation of coarse deep learning prediction maps for the segmentation of fine structures in immunofluorescence images. In Medical Imaging 2018: Digital Pathology, volume 10581, page 105810P. International Society for Optics and Photonics, 2018.

[72] Olaf Ronneberger, Philipp Fischer, and Thomas Brox. U-net: Convolutional networks for biomedical image segmentation. In International Conference on Medical image computing and computer-assisted intervention, pages 234–241. Springer, 2015.

[73] Brian D Ripley. Modelling spatial patterns. Journal of the Royal Statistical Society: Series B (Methodological), 39(2):172–192, 1977.

[74] J Besag. Contribution to the discussion on dr ripley’s paper. JR Stat. Soc., 39:193–195, 1977.

[75] Julienne L Carstens, Pedro Correa De Sampaio, Dalu Yang, Souptik Barua, Huamin Wang, Arvind Rao, James P Allison, Valerie S LeBleu, and Raghu Kalluri. Spatial computation of intratumoral t cells correlates with survival of patients with pancreatic cancer. Nature communications, 8:15095, 2017.

[76] KM. Leung, M. R. Elashoff, and A. A. Afifi. Censoring issues in survival analysis. Annual Review of Public Health, 18(1):83–104, May 1997.

[77] Niko Kemi, Maarit Eskuri, and Joonas H Kauppila. Tumour-stroma ratio and 5-year mortality in gastric adenocarcinoma: a systematic review and meta-analysis. Scientific reports, 9(1):1–6, 2019.

[78] AP Noon, PC Albertsen, F Thomas, DJ Rosario, and JWF Catto. Competing mortality in patients diagnosed with bladder cancer: evidence of undertreatment in the elderly and female patients. British journal of cancer, 108(7):1534–1540, 2013.

[79] J. Bergstra, D. Yamins, and D. D. Cox. Making a science of model search: Hyperparameter optimization in hundreds of dimensions for vision architectures. In Proceedings of the 30th International Conference on International Conference on Machine Learning, volume 28, pages I-115–I-123, 2013.

[80] J. Bergstra and Y. Bengio. Random search for hyper-parameter optimization. Journal of Machine Learning Research, 13:281–305, 2012.

[81] D. H. Wolpert and W. G. Macready. No free lunch theorems for optimization. Trans. Evol. Comp, 1(1):67–82, 1997.

