## Supplementary Information for "Identification of Immunological Features Enables Survival Prediction of Muscle-Invasive Bladder Cancer Patients Using Machine Learning"

February 24, 2020

### Supplementary Information

Table S1: The features that contribute to a good and bad prognosis according to the LR and the LSVM. L(x,y,r): the L function value of $y$ in respect to $x$ for distance $r$.

| Classifiers | Bad Prognosis | Good Prognosis |
|---|---|---|
| **LR** | Density of TB frontin/core | Density of $CD8^+$ frontin/frontout/core<br>Density of $CD3^+$ frontout/core<br>Density of $CD68^+$ frontin/frontout<br>L(TB, $CD3^+$, 20) |
| **LSVM** | Density of TB frontin/core | Density of $CD8^+$ frontin/frontout/core<br>Density of $CD3^+$ frontin/frontout/core<br>Density of $CD68^+$ frontin/frontout/core<br>Number of $CD3^+$ frontin<br>Number of $PDL1^+CD163^-CD68^+$ frontout<br>Number of $PDL1^+CD163^+CD68^+$ frontout |

*Equal contribution

Table S2: The most important features for estimating patient prognosis by the DT and the RF. L(x,y,r): the L function value of $y$ in respect to $x$ for distance $r$.

| Classifiers | Important Features |
|---|---|
| **DT** | Number of PD-L1$^+$ frontout |
| | Density of CD163$^+$ frontout |
| | Density of PD-L1$^+$CK$^+$ core |
| | Number of TB frontout |
| | Density of CD3$^+$ frontout |
| | Number of CD68$^+$ frontout |
| | Density of CD68$^+$ frontin/frontout |
| **RF** | Number of CD68$^+$ frontout |
| | Density of CD68$^+$ frontin/frontout/core |
| | Number of CD3$^+$ core |
| | Density of CD3$^+$ frontout |
| | Number of CD8$^+$ frontout/core |
| | Density of CD8$^+$ frontout/core |
| | Number of TB core |
| | Density of PD-L1$^+$ frontout |
| | Density of PD-L1$^+$CK$^+$ core |
| | Density of PD-L1$^+$CK$^-$ frontin/frontout |
| | Number of CD163$^+$CD68$^+$ frontout |
| | Number of NucleiCK$^+$ frontin |
| | TNM IIIA |
| | TNM IV |
| | L(TB, CD3$^+$, 20) |
| | L(TB, CD8$^+$, 20) |
| | L(TB, PD-L1$^+$, 20) |
| | L(TB, CD8$^+$, 50) |
| | L(CD163$^+$, PD-L1$^+$, 150) |

Table S3: Results for algorithm selection from the nested cross validation on the training set with AUROC as the performance metric. For each feature space, the best ML classifier is indicated in bold. Amongst the best classifiers of each feature space (in bold), our ensemble model uses those with a marked difference in performance (**).

| Feature Space | LR | KNN | LSVM | RSVM | DT | RF |
|---|---|---|---|---|---|---|
| **Image** | $69.8 \pm 13.3$ | $61.8 \pm 5.7$ | $**$ **72.8 ± 0.3** | $60.8 \pm 8.2$ | $63.3 \pm 3.3$ | $57.8 \pm 3.2$ |
| **Clinical** | **58.1 ± 0.1** | $59.8 \pm 7.2$ | $55.9 \pm 5.9$ | $48.6 \pm 5.6$ | $50.1 \pm 2.4$ | $56.6 \pm 9.1$ |
| **Spatial** | $46.8 \pm 3.3$ | $46.2 \pm 4.7$ | $42.7 \pm 1.8$ | $38.0 \pm 0.0$ | **49.2 ± 3.3** | $43.7 \pm 13.0$ |
| **Image & clinical** | $62.0 \pm 0.6$ | $55.3 \pm 4.4$ | $56.5 \pm 8.5$ | $50.1 \pm 11.5$ | $**$ **68.8 ± 0.8** | $70.4 \pm 7.9$ |
| **Image & spatial** | $**$ **70.2 ± 14.7** | $59.9 \pm 2.4$ | $49.0 \pm 2.5$ | $58.6 \pm 11.9$ | $51.0 \pm 1.5$ | $64.1 \pm 7.6$ |
| **Clinical & spatial** | $44.3 \pm 9.8$ | $49.4 \pm 3.2$ | **57.9 ± 2.4** | $47.1 \pm 4.4$ | $44.8 \pm 2.7$ | $46.6 \pm 15.4$ |
| **Image & clinical & spatial** | $56.3 \pm 0.2$ | $55.0 \pm 4.0$ | $60.2 \pm 8.2$ | $61.4 \pm 4.6$ | $66.6 \pm 9.1$ | $**$ **67.3 ± 5.8** |

**(a)**

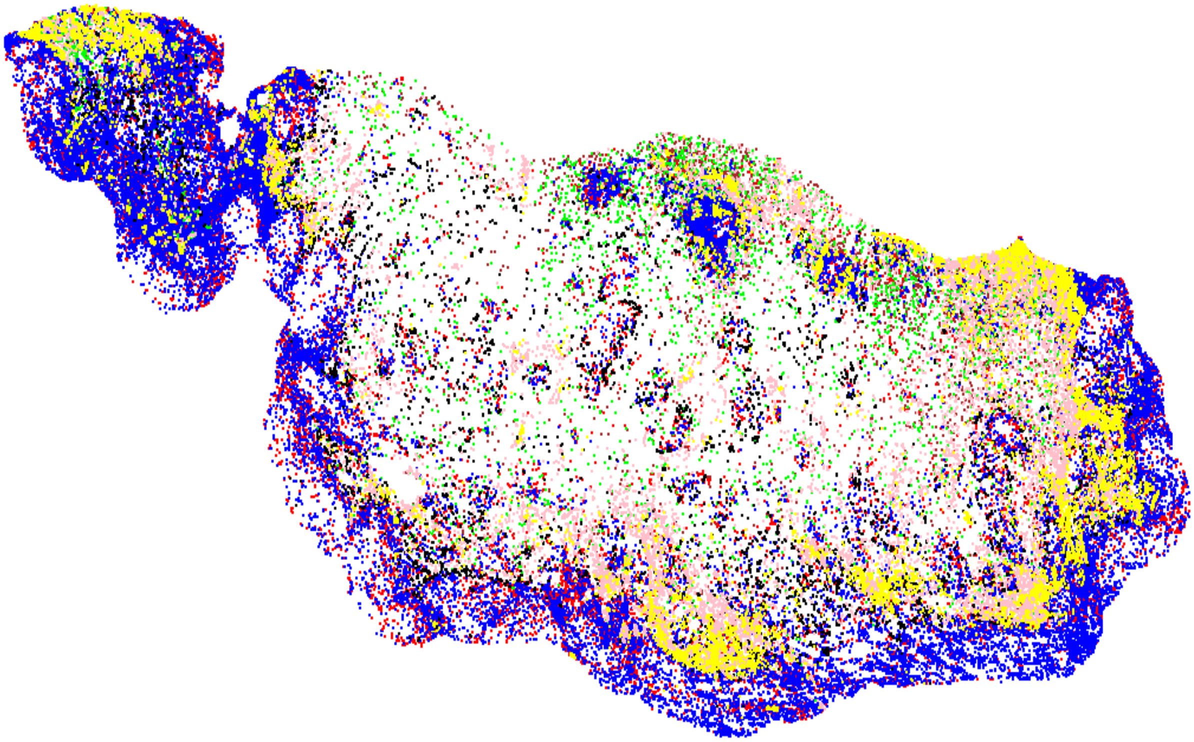

**(b)**

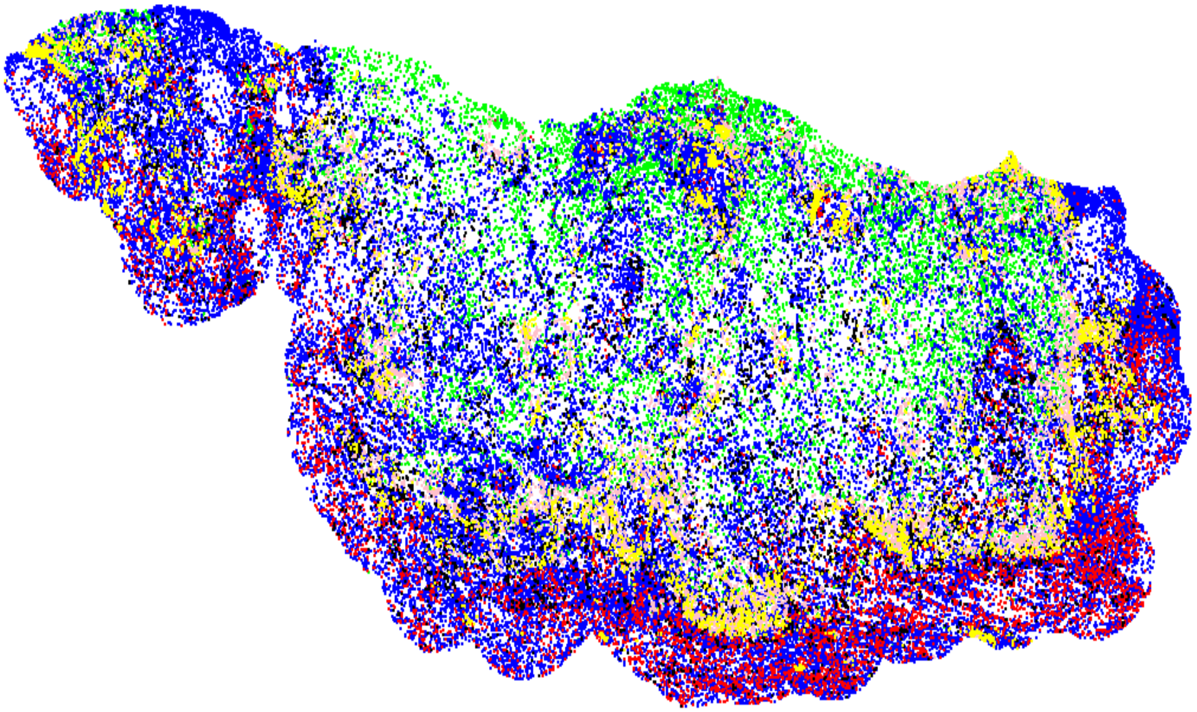

Figure S1: Nuclei localization of (a) TILs (*$CD3^+CK^+$: Brown, $CD3^+CK^-$: Red, $CD8^+CK^+$: Green, $CD8^+CK^-$: Blue, TB: black, $PD\text{-}L1^+CK^+$: Pink, $PD\text{-}L1^+CK^-$: Yellow*) and (b) TAMs (*$CD163^+CK^+$: Brown, $CD163^+CK^-$: Red, $CD68^+CK^+$: Green, $CD68^+CK^-$: Blue, TB: black, $PD\text{-}L1^+CK^+$: Pink, $PD\text{-}L1^+CK^-$: Yellow*) across the WSI.

**(a)**

**(b)**

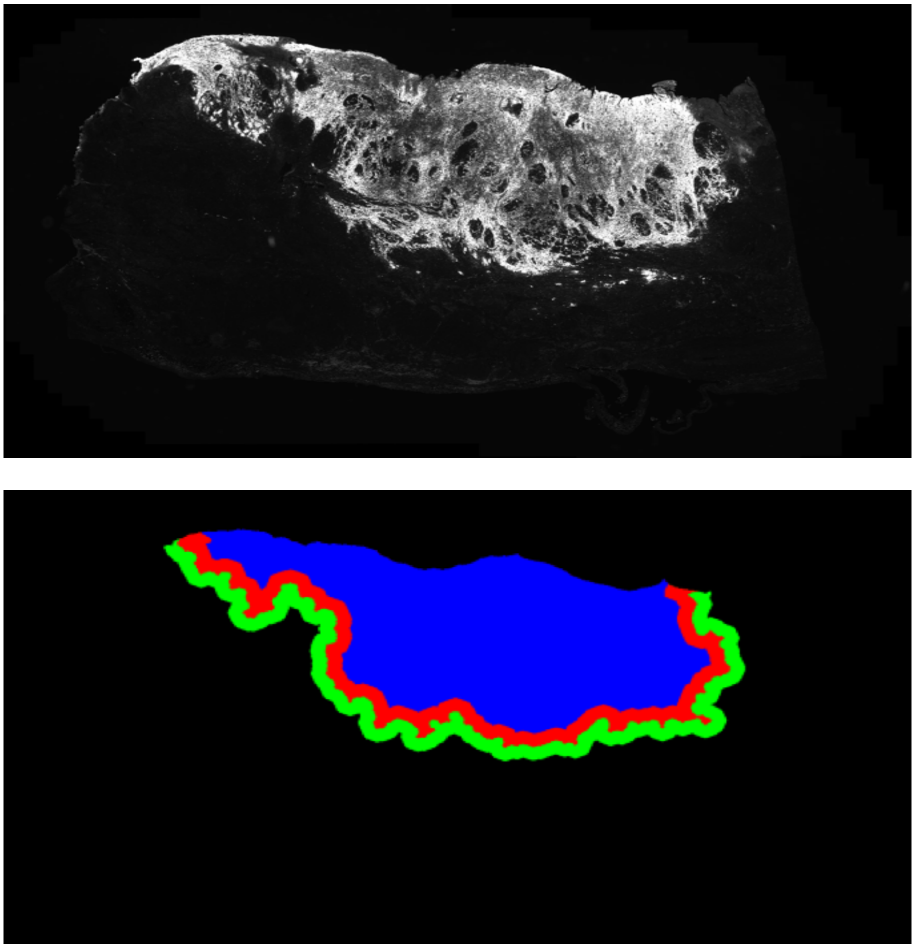

Figure S2: (a) Whole slide immunofluorescence image based on the PanCK channel, (b) Segmentation of the corresponding tissue (a) into tumour core (*blue*), invasive frontin (*red*) and frontout (*green*) using the PanCK channel.

**(a)** **(b)**

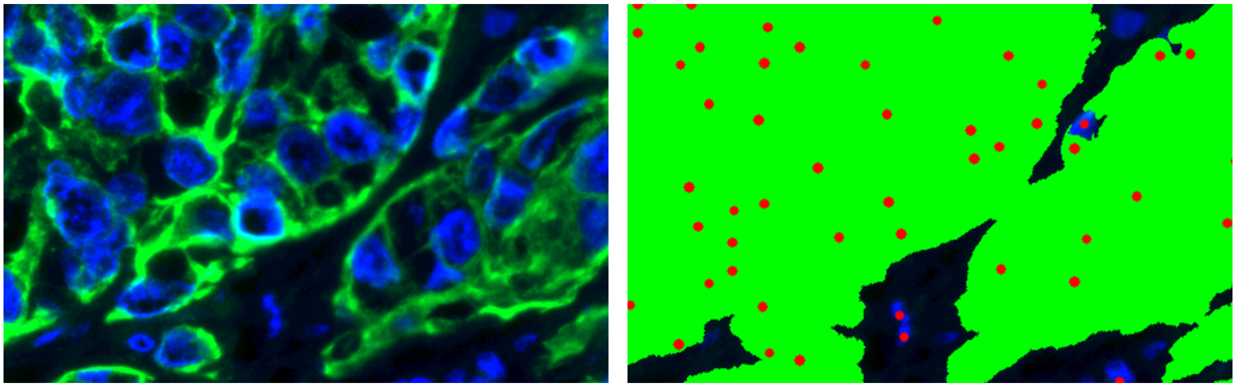

Figure S3: (a) Immunofluorescence region visualised using the PanCK and Hoechst channels, (b) The corresponding epithelium segmentation mask of (a) along with the detected cell nuclei.

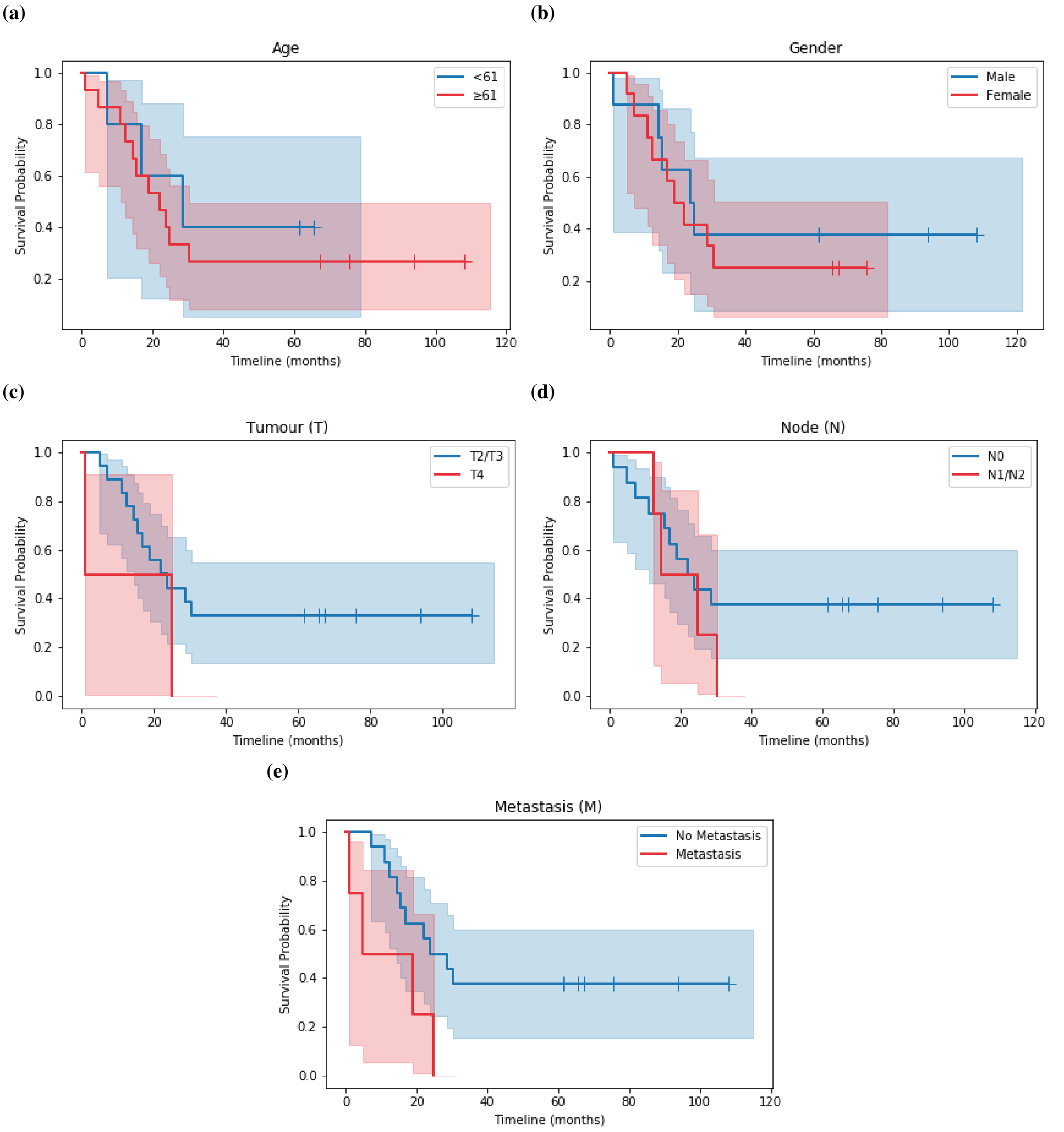


Figure S4: Kaplan-Meier curves of (a) Age ($p$ value $= 0.57$, $N_{<61} = 5$ & $N_{\geq 61} = 15$), (b) Gender ($p$ value $= 0.62$, $N_{Female} = 12$ & $N_{Male} = 8$), (c) Tumour ($p$ value $= 0.25$, $N_{T2/T3} = 10$ & $N_{T4} = 2$), (d) Node ($p$ value $= 0.36$, $N_{N0} = 16$ & $N_{N1/N2} = 4$), and (e) Metastasis ($p$ value $= 0.04$, $N_{No} = 16$ & $N_{Yes} = 4$).

**(a)**

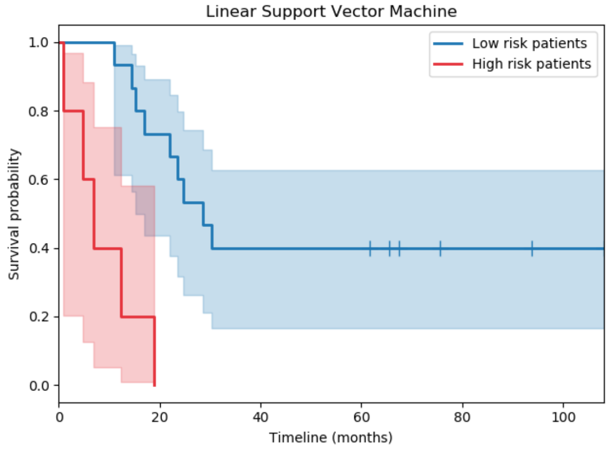
Linear Support Vector Machine
Low risk patients
High risk patients
Survival probability
Timeline (months)
0.0
0.2
0.4
0.6
0.8
1.0
0
20
40
60
80
100

**(b)**

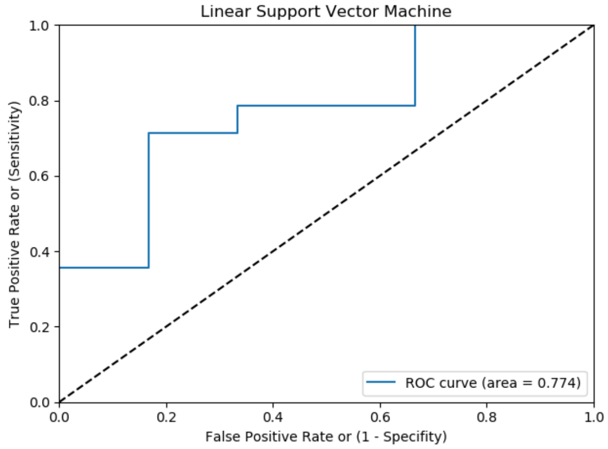
Linear Support Vector Machine
True Positive Rate or (Sensitivity)
False Positive Rate or (1 - Specifity)
ROC curve (area = 0.774)
0.0
0.2
0.4
0.6
0.8
1.0

**(c)**

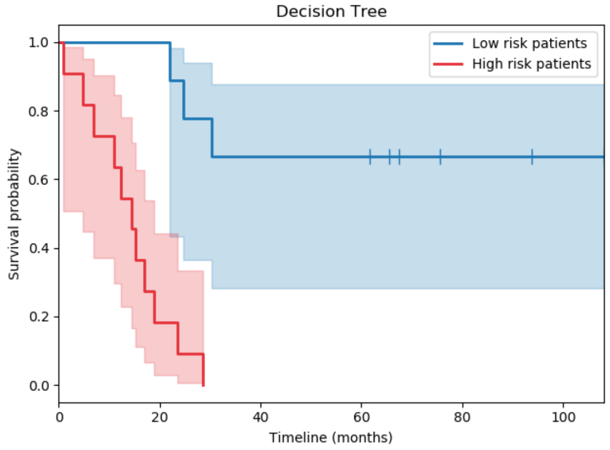
Decision Tree
Low risk patients
High risk patients
Survival probability
Timeline (months)
0.0
0.2
0.4
0.6
0.8
1.0
0
20
40
60
80
100

**(d)**

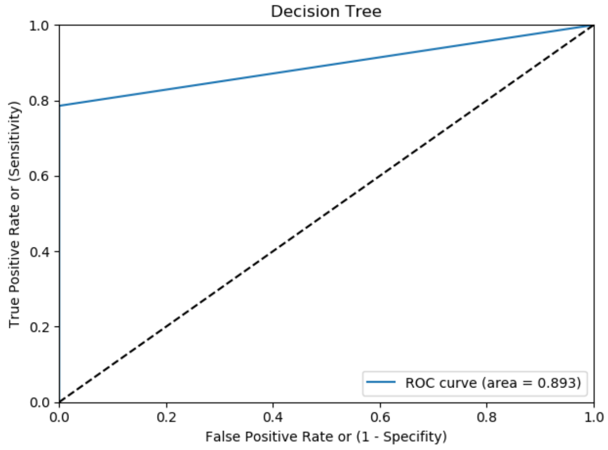
Decision Tree
True Positive Rate or (Sensitivity)
False Positive Rate or (1 - Specifity)
ROC curve (area = 0.893)
0.0
0.2
0.4
0.6
0.8
1.0

**(e)**

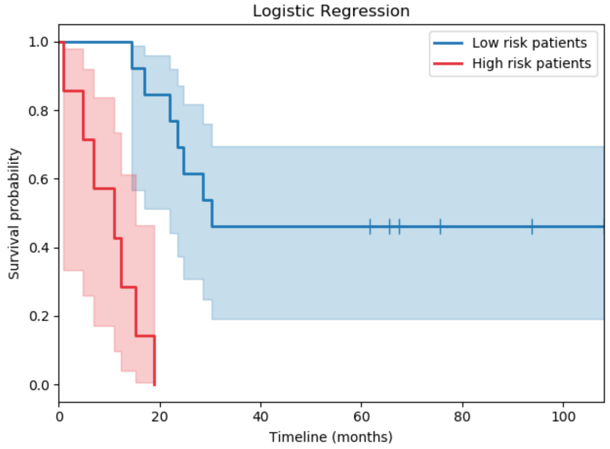
Logistic Regression
Low risk patients
High risk patients
Survival probability
Timeline (months)
0.0
0.2
0.4
0.6
0.8
1.0
0
20
40
60
80
100

**(f)**

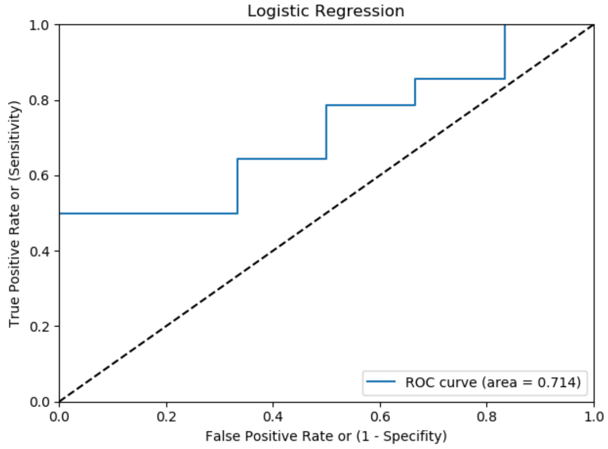
Logistic Regression
True Positive Rate or (Sensitivity)
False Positive Rate or (1 - Specifity)
ROC curve (area = 0.714)
0.0
0.2
0.4
0.6
0.8
1.0

(g) (h)

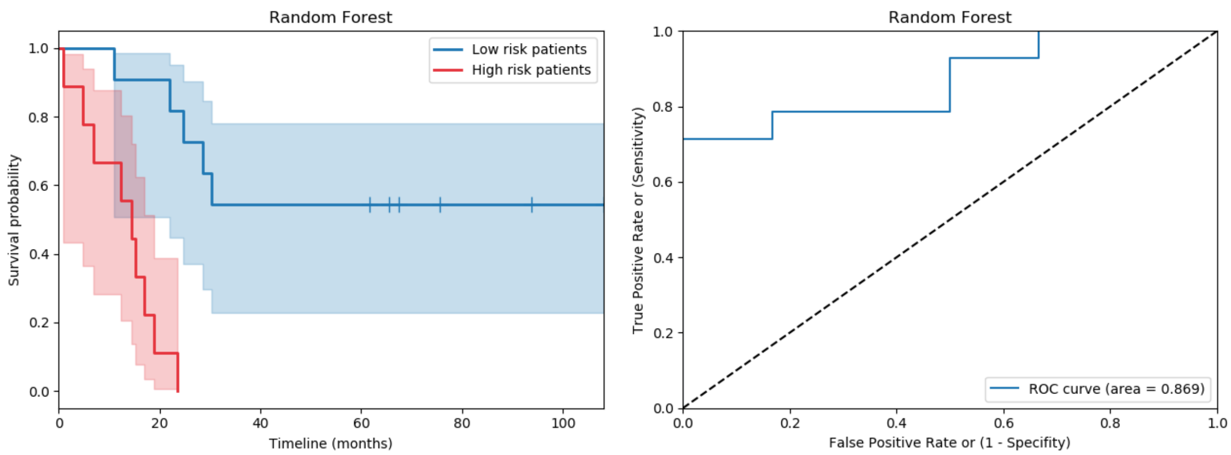


Figure S5: Kaplan-Meier and ROC curves on the testing set for each of the ML classifiers used in the ensemble model. (a–b) Image features ($p$ value $= 6e-05$, $N_{LowRisk} = 15$ & $N_{HighRisk} = 5$), (c–d) image and clinical features ($p$ value $= 1e-04$, $N_{LowRisk} = 9$ & $N_{HighRisk} = 11$), (e–f) image and spatial features ($p$ value $= 5e-05$, $N_{LowRisk} = 13$ & $N_{HighRisk} = 7$), (g–h) image, clinical, and spatial features ($p$ value $= 8e-06$, $N_{LowRisk} = 11$ & $N_{HighRisk} = 9$).

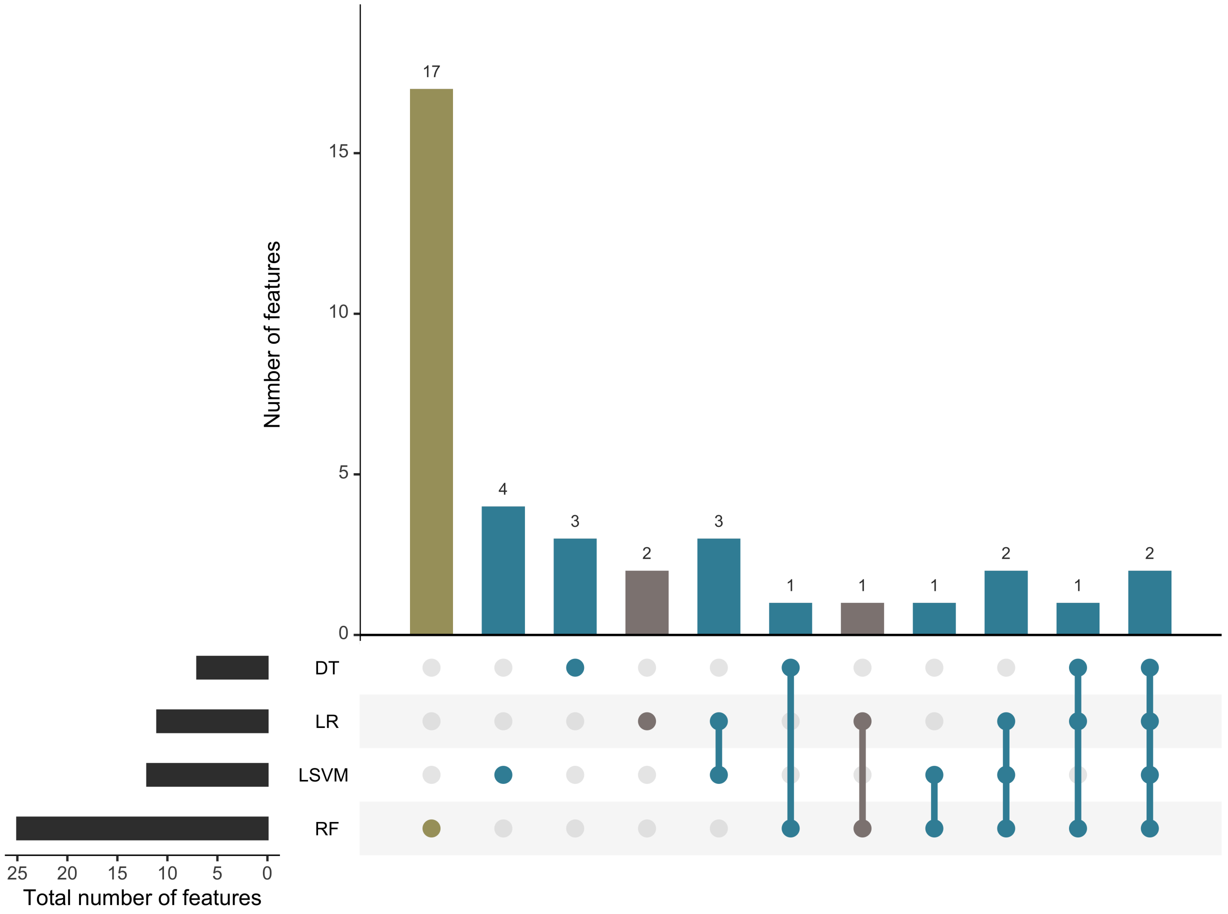


Figure S6: Intersecting features between the various submodels of our ensemble model. Gray denotes image and spatial features, blue denotes image features, and mustard denotes image, clinical, and spatial features.

Table S4: The search space of each classifier based on predefined distributions over its hyperparameters.

| Classifier | Hyperparameter | Distribution | Values |
|---|---|---|---|
| LSVM | C | Log-uniform | $[\ln(1e{-}5),\ \ln(1e2)]$ |
| | Class weight | Categorical | [Balanced, None] |
| RSVM | C | Log-uniform | $[\ln(1e{-}5),\ \ln(1e2)]$ |
| | Gamma | Log-uniform | $1/\text{max\_features} * [\ln(1e{-}3), \ln(1e3)]$ |
| | Class weight | Categorical | [Balanced, None] |
| LR | Type of penalty | Categorical | [$l1$, $l2$, Elastic net, None] |
| | C | Log-uniform | $[\ln(1e{-}5),\ \ln(1e2)]$ |
| | L1 ratio | Uniform | $[0,\ 1]$ |
| | Class weight | Categorical | [Balanced, None] |
| DT | Criterion | Categorical | [Gini, Entropy] |
| | Maximum features | Uniform integer | [1, max_features] |
| | Maximum depth | Categorical | $[1, 15]$ or None |
| | Class weight | Categorical | [Balanced, None] |
| RF | Number of trees | Log-uniform integer | $[10,\ 1000]$ |
| | Criterion | Categorical | [Gini, Entropy] |
| | Maximum features | Uniform integer | [1, max_features] |
| | Maximum depth | Categorical | [$2, 3, 4$, None] |
| | Bootstrap | Categorical | [True, False] |
| | Class weight | Categorical | [Balanced, None] |
| KNN | K | Log-uniform integer | [1, 20] |
| | Metric | Categorical | [Balanced, None] |
| | P | Uniform integer | [1, 6] |

Table S5: Immunofluorescence primary antibodies.

| Antibody | Supplier | Catalogue number | Species | Dilution |
|---|---|---|---|---|
| CD3 | Agilent Technologies | A0452 | Rabbit-Polyclonal | 1:400 |
| CD8 | Agilent Technologies | M7103-Clone C8/144B | Mouse-Polyclonal | 1:200 |
| CD68 | Cell Signaling | D4B9C | Rabbit-Monoclonal | 1:3000 |
| CD163 | Cell Marque | MRQ-26 | Mouse-Monoclonal | 1:3000 |
| PD-L1 | Cell Signaling | 13684S | Rabbit-Monoclonal | 1:100 |
| PanCK | Agilent Technologies | Z0622 | Rabbit-Polyclonal | 1:100 |

Table S6: Detection and visualization reagents for target proteins.

| Dye | Supplier | Catalogue number | Dilution |
|---|---|---|---|
| FITC | Perkin Elmer | NEL741B001KT | 1:100 |
| CY3 | Perkin Elmer | NEL744B001KT | 1:100 |
| CY5 | Perkin Elmer | NEL745B001KT | 1:100 |
| Alexa Fluor 750 | ThermoFisher Scientific | S21384 | 1:50 |

Table S7: Summary of each immunofluorescence target. Visualization and imaging acquisition information.

| Target | Dye name | Excitation wavelength (nm) | Emission wavelength (nm) | Exposure time (ms) |
|---|---|---|---|---|
| Nucleic acid (DNA) | Hoechst | 353 | 465 | 10 |
| CD3 | FITC | 495 | 519 | 10 |
| CD8 | CY3 | 548 | 561 | 10 |
| PD-L1 | CY5 | 650 | 673 | 10 |
| PanCK | CY7 | 747 | 773 | 600 |

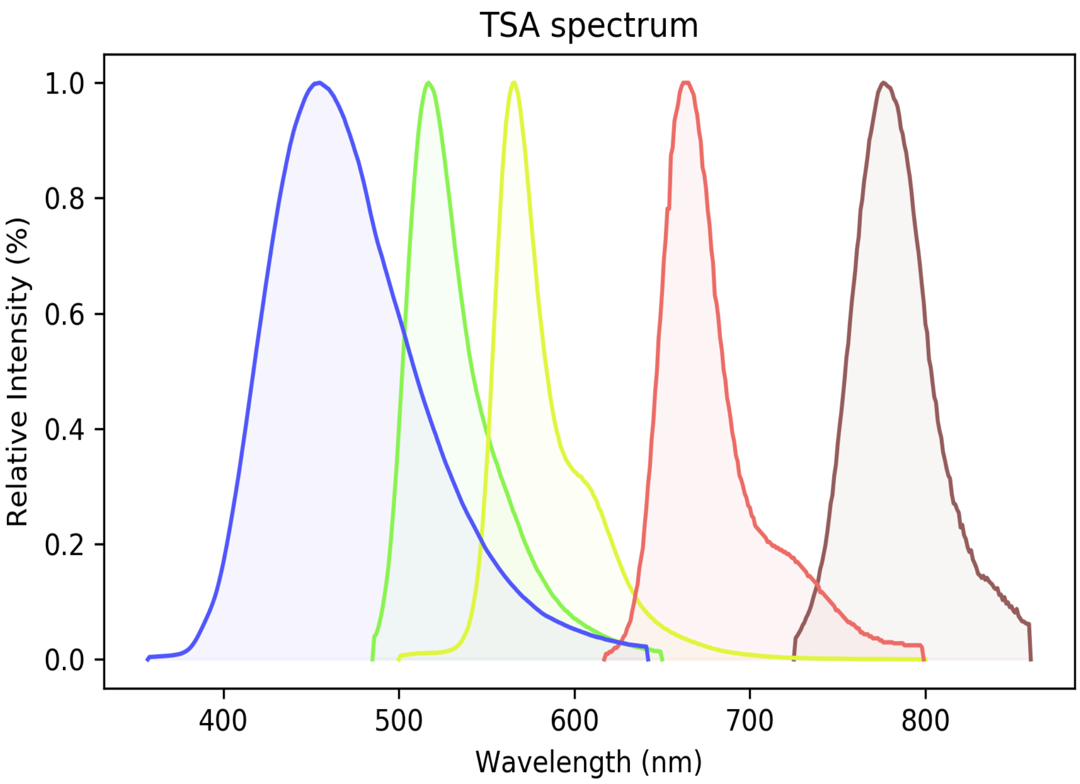


Figure S7: Tyramide signal amplification spectra for antibody visualisation. *Hoechst: Blue, FITC: Green, CY3: Yellow, CY5: Soft red, Alexa Fluor 750 (CY7): Dark red* .

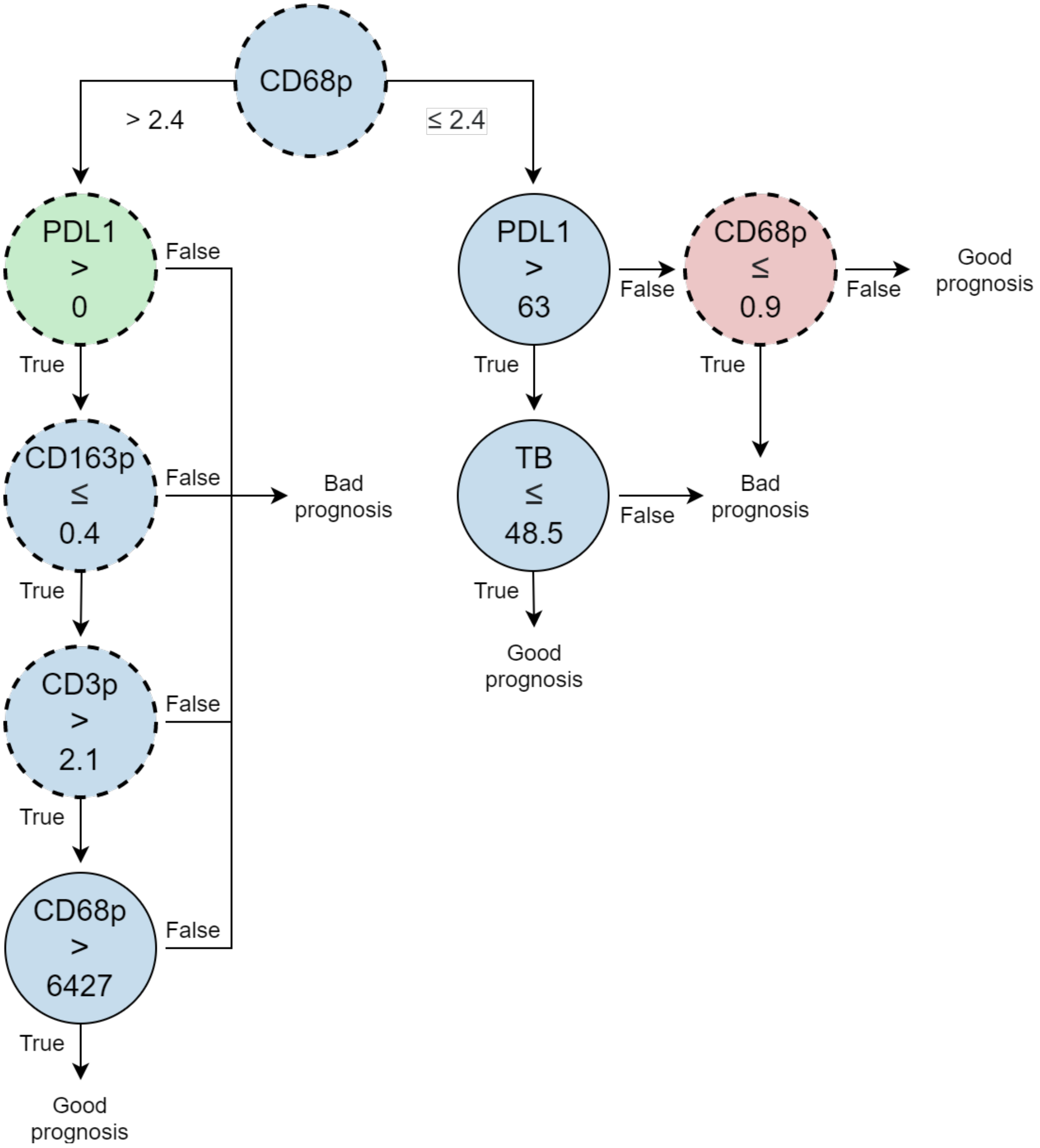


Figure S8: Diagram of the DT model employed by the ensemble model. *Solid-line circles: number of cells, dashed-line circles: density of cells. Tumour core in green, frontin in red and frontout in blue.*

Table S8: Pairwise comparison between TNM staging groups. Stages IIIA and IIIB were merged due to having the largest $p$ value.

| TNM | II | IIIA | IIIB |
|---|---|---|---|
| IIIA | 0.33792 | - | - |
| IIIB | 0.50382 | 0.93012 | - |
| IV | 0.00087 | 0.00317 | 0.07361 |

Table S9: Pairwise comparison between TNM staging groups. Stages II and IIIA-IIIB were merged due to having the largest $p$ value.

| TNM | II | IIIA-IIIB |
|---|---|---|
| IIIA-IIIB | 0.21458 | - |
| IV | 0.00034 | 0.00034 |

Table S10: Pairwise comparisons using log-rank test between TNM staging groups. Since the $p$ value was significant, the two stages were not merged.

| TNM | II-IIIA-IIIB |
|---|---|
| IV | 1.6e-06 |
